# A new highly specific and soluble protease for precise removal of N-terminal purification tags

**DOI:** 10.1101/2024.09.17.613541

**Authors:** Raef Shams, Lynne Regan

**Author notes:** **Correspondence:** Lynne Regan, Roger Land Building, Alexander Crum Brown Road, King’s Buildings, Edinburgh EH9 3FF. **Address**: Roger Land Building, Alexander Crum Brown Road, King’s Buildings, Edinburgh EH9 3FF, United Kingdom. **Emails**.

## Abstract

The affinity purification tags have significantly enhanced the ease of purification of many proteins. After purification, it is necessary to remove the affinity purification tag for many applications. Several specific proteases have been developed for tag removal, but none yet has both, a desirable cleavage specificity and high solubility. Here we describe the design and characterization of a novel protease, which we have named Con1, which exhibits desirable cleavage specificity and high solubility. We present a detailed analysis of the kinetic properties of Con1, along with demonstrating its ability to remove purification tags from different proteins.

## 1. Introduction

Affinity purification of recombinant proteins is a powerful tool [1–4]. The use of protein purification tags – such as polyhistidine (His_n_, where n is typically 6), glutathione S transferase (GST) or maltose binding protein (MBP) has greatly simplified the purification of recombinantly expressed proteins [1–3]. The fusion of a purification tag with a protein of interest (POI) means that a standard purification method can be used for various POIs [4]. The fusion tag is often placed at the N-terminus of the POI so it can be enzymatically removed after purification [5, 6]. Ideally, the affinity tags must be removed from the POI, and this is often accomplished by incorporating a protease cleavage site between the affinity tag and the POI [4]. Several enzymes have been developed for such purification tag removal, one of the most widely used is Tobacco Etch Virus nuclear-inclusion-A endopeptidase (TEV protease), which is favoured because of its high specificity [7]. TEV protease has a recognition sequence ENLYFQ-G/S, where (-) represents the site of cleavage [8]. The naming of the residues follows the convention P_6_-P_5_-P_4_-P_3_-P_2_-P_1_-P_1_’. For TEV, the preferred residue at the P_1_’ position is Gly or Ser, which means that after cleavage a Gly or Ser residue is added to the N-terminus of the POI [9]. For many laboratory applications, the addition of an extra residue to the N-terminus of the POI is not problematic. If a recombinant POI is to be a biotherapeutic, however, it must be closely similar to the natural source protein ‘the reference drug’, for which there is already regulatory approval. Thus, the addition of extra amino acids to the N-terminus is not desirable. Although TEV protease is widely used, it is well known to exhibit low solubility that diminishes the purification yield, high concentration storage and use [10, 11]. To date, great effort has been made to improve TEV protease solubility, however, the solubility is still low [10, 12]. With these attributes as motivation, we sought a protease with high substrate specificity, with little restriction on the identity of the P_1_’ residue, and with good solubility.

## 2. Materials and Methods

### 2.1. Computational work

#### 2.1.1. FASTA search for TEV-like proteases

The full-length sequence of the TEV protease (1-242), also known as Nuclear Inclusion protein a (NIa), was taken from the UniProt database (UniProt code of the poly-protein origin: <u>P04157</u>; Genome position in TEV virus: 2038-2279) [13]. We performed a sequence similarity search of this sequence against the UniProt Knowledgebase and UniProtKB/ Swiss-Prot isoforms databases using the <u>FASTA</u> suite of programmes [14]. We shortlisted the results by excluding variants with ≤50% (which is the desired threshold) and ≥90% (which are mutant TEV protease versions) identity to the TEV protease sequence.

#### 2.1.2. Intrinsic solubility prediction

We used the CamSol, web server to predict protein solubility based on the amino acid sequence [15–17]. Scores >0 indicate soluble proteins, whereas scores of <0 indicate poorly soluble proteins. The higher the score, the more soluble a protein is predicted to be.

#### 2.1.3. Alignment and consensus

We aligned the full-length sequence of the selected proteases (TEV, TVMV, TuMV_Q, TuMV_J, PPV, OMV, LMVE and LMV0) using the Clustal Omega tool using the default settings [14]. The alignment was then exported and analyzed by the Jalview 2.11.2.6 software [18]. The alignment was coloured by the Clustal scheme with >75% identity threshold. The software automatically generates the consensus sequence as a percentage (>40% conservation) of the modal residue per column. A clear consensus backbone that comprised 207 conserved residues was identified. At this point, the above-mentioned substrate binding positions were retained in the consensus designs to the equivalent TuMV_J residues. These are: N30, A170, I176, S214, E219, and S220. The remaining non-conserved 36 positions were filled by choosing residues of the original sequences based on their hydrophilicity (Cys residues were excluded). These combinations allowed us to build 10 different full-length variants (Con1-Con10). Finally, C-terminal (235-243) was subjected to deletion to improve solubility since most TEV protease studies were performed using C-terminal truncated versions [8, 11, 19].

### 2.2. Experimental works

#### 2.2.1. Plasmid construction

##### A- Proteases

We obtained synthetic genes encoding 7xHis tagged TuMV_J and Con1 (IDT, Belgium, Supplementary notes). These were cloned by PCR to a pMAL expression vectors using AQUA cloning [20]. Following transformation into the *E. coli* strain Top 10, the desired plasmids were identified by DNA sequencing. See supplementary materials for genes and primer sequences. C-terminal deletions of Con1 were performed using PCR with primers designed to delete the nucleotides corresponding to amino acids 228-234 or 222-234 (Supplementary Table 1).

##### B- FRET substrates

The synthetic genes encoding CFP-linker-cleavage-site-linker-YFP (IDT, Belgium, Supplementary notes) were cloned by digestion/ligation into the backbone of a pET28 expression vector. The inserts and a vector were digested by BamHI/BsrGI enzymes and ligated by T4 ligase (NEB). Correct inserts were verified by DNA sequencing.

##### C- His-tagged POI

We designed constructs encoding the N-terminal His tag separated from the POI by the P6-P1 residues of the Con1 cleavage site. The N-terminal residue of the POI contributes to the P1’ position of the cleavage site. The correct clones were identified by DNA sequencing.

#### 2.2.2. Protein expression

Proteins were expressed using standard protocols. Briefly, colonies of BL21(DE3) containing the desired construct were inoculated into 5 ml of LB plus antibiotic and grown overnight at 37°C. The next day, the overnight cultures were added to LB plus antibiotic media and grown with shaking at 37°C until the OD_600_ was 0.6. IPTG was then added to a final concentration of 1 mM, the temperature was dropped to 20°C, and the incubation continued overnight. The following day, cells were harvested by centrifugation and stored at −80°C.

#### 2.2.3. Protein purification

##### A- Con1, TuMV_J and TEV proteases

Cell paste (from 100 ml culture) was re-suspended in 10 mL buffer A (50 mM Tris-HCl, pH 8.0, 100 mM NaCl, and 5% (v/v) glycerol) supplemented with protease inhibitor cocktail tablets (cOmpleteTM, Roche). The suspensions were kept on ice, and cells were disrupted by sonication. After sonication, the solution was centrifuged to remove insoluble material, and the pellet was discarded. Supernatants were applied to a 5ml Ni-NTA agarose column pre-equilibrated in buffer A. After sample loading, the resin was washed with buffer A plus 20 mM imidazole. Finally, proteins were eluted by buffer A with an imidazole concentration from 75 to 500mM. The fractions were analysed by SDS–PAGE, pooled, buffer exchanged into buffer A and concentrated using 3 kDa MW-CO Amicon (Merck-Millipore).

##### B- FRET substrates, His-POI and proteases (Con1, ^Δ228-234^Con1 and ^Δ222-234^Con1)

Cell lysates were prepared as described above. The supernatant was applied to a HisTrap Excel column (Cytiva) equilibrated in buffer A and eluted by a gradient of 0–1M Imidazole, in buffer A. The peak fractions were analysed by SDS–PAGE, pooled, buffer exchanged and concentrated using a 3 kDa MW-CO Amicon, then stored at −80°C.

FRET substrates and proteases were additionally purified by gel filtration using a Superdex-75 column, equilibrated, and run-in buffer A. After checking the purity on SDS–PAGE, appropriate fractions were pooled, concentrated using a 3 kDa MW-CO proper name (Amicon), and then stored at −80°C.

#### 2.2.4. Protease solubility

The solubility of TEV and Con1 proteases was studied by concentrating proteins and collecting aliquots over time as previously described [10]. The 5 ml aliquots of 1.5 mg/ml proteins were spun down at high speed to pellet any aggregates before the experiment began. Protein concentrations were then measured by the Nano-drop spectrophotometer using the extinction coefficients of 31970 and 34950 M-1cm-1 for TEV (MW= 28560.45 Da) and Con1 (MW=27660.22 Da), respectively. The proteins were loaded onto a 5-ml 3 kDa MW-CO Amicon and centrifuged at 4000 rpm at 4°C to concentrate until the aggregation was observed.

High concentrations of TEV (15 mg/ml) or Con1 (30 mg/ml) proteases were incubated overnight at room temperature, aliquots were collected over time, and optical density (OD) at 400 nm was measured.

#### 2.2.5. Proteolytic activity

##### A- Substrate specificity

We assessed the specificity of Con1, TEV and TuMV_J for the cleavage sites EAVYHQ-S and ENLYFQ-G. Protease (1 µM) and substrate (2 µM) were incubated in Buffer A at room temperature. The mixtures were excited at 430 nm, and fluorescence emission intensity at 480 nm and 520 nm were measured every minute over 2 hours. The amount of cleavage was determined from the change in FRET ratio, calculated by the intensity at 480 nm (donor) divided by the intensity at 520 nm (acceptor), after blank subtraction. Cleavage is a percentage of the endpoint after saturation. All the experiments were performed in duplicate.

To screen the P1’ specificity of Con1, FRET substrates with different cleavage sequences between the donor and acceptor fluorescent proteins were prepared: EAVYHQ-X (X= S, D, M or P). A 0.5 µM Con1 was mixed with 5 µM substrate in Buffer A and the fluorescence was measured over 5 hours at room temperature.

##### B- Kinetics of Con1 Vs. TuMV_J

Kinetic parameters for the proteases Con1 and TuMV_J were determined by incubating 0.4 µM enzyme with the substrates at range of concentrations (1-24 µM) in Buffer A. Different FRET substrates EAVYHQ-X (X = Ala, Ser, Met or Asp) between the FRET donor (CFP) and acceptor (YFP) proteins were used to assess the influence of the identity of the P1’ position.

##### C- Kinetics of Con1 at different pH

Con1 and substrate fractions were pre-incubated at different pHs of 0.1M citrate, pH 6.0; 0.1M Tris-HCl, pH 7.0; 0.1M Tris-HCl, pH 8.0 or 0.1M Tris-HCl, pH 9.0. The reaction mixtures containing 0.4 µM Con1 incubated with an increasing concentration of the FRET substrate incorporating the EAVYHQ-M site (0.3125, 0.625, 1.25, 2.5, 5, 7.5, 10 and12.5 µM) at different pH.

##### D- Kinetics of Con1 Vs. C-terminal truncated versions

Kinetic parameters for Con1, ^Δ228-234^Con1 and ^Δ222-234^Con1 (0.4 µM) against FRET substrate incorporating EAVYHQ-M site (0.625-25 µM) were determined at 50 mM Tris-HCl, pH 7.0, 200 mM NaCl buffer.

For the kinetic studies, the data were collected and analysed as previously described [21]. The full-scale range (FSR) was calculated for each wavelength by subtracting the minimum emission (Em^min^; which is at time zero intensity at 480 nm and the end-point intensity at 520 nm) from the maximum emission (Em^max^; which is at end-point intensity at 480 nm and the time zero intensity at 520 nm by the following equations:

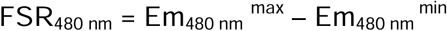

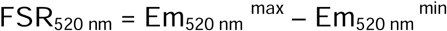

Then, the emission values were normalised by subtracting the emission at time zero by the following equations:

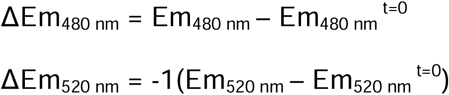

Then, the cleavage percentages (C^%^) were calculated by dividing the normalised values by the FSR values by the following equations:

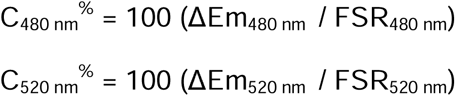

The cleaved product concentration [P] was calculated as a percentage of the total substrate concentration at time zero by the following equation:

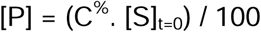

After that, the product concentrations (on the Y axis) were plotted vs. time (on the X axis) and the curves were fitted by the exponential plateau equation on GraphPad Prism software that properly fits frequency changes versus time:

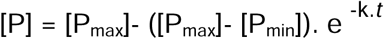

Where P_max_; is the maximum product concentration, P_min_; is the minimum product concentration, k; is the rate constant, and *t* is the time. From the calculated rate constants, the reaction rates (V) were calculated by the following equation:

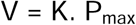

Finally, the reaction rates (Y-axis) were plotted vs. substrate concentration (X-axis) and the curves were fitted by the Michaelis-Menten equation on GraphPad Prism software:

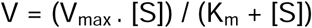

Where V is the reaction rate; V_max_ is the maximum velocity; [S] is the substrate concentration; and K_m_ is the Michaelis constant. The V_max_ and K_m_ were obtained from the non-linear regression and the K_cat_ was calculated by the following equation:

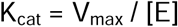

Where Kcat is the catalytic turnover number and [E] is the total enzyme concentration.

#### 2.2.6. pH dependency of Con1 activity

##### A. FRET assay

To evaluate the pH dependence of the Con1 activity, we performed the FRET cleavage assay in buffers of different pH: 0.1 M Sodium citrate (pH 4, 5 or 6), 0.1 M Tris-HCl (pH 7, 8 or 9) and 0.1 M sodium borate (pH 10) [22]. The Con1 protease (0.5 µM) and the FRET substrate (2.5 µM) were pre-incubated in the different pH buffers at room temperature for 10 min. The reaction was initiated by mixing protease with substrate, and incubation continued for a further 2 hours at room temperature. The end-point fluorescence was measured at 480 and 520 nm and the amount of cleavage was calculated from the change in FRET.

##### B. SDS-PAGE experiments

Con1 and His7-(EAVYHQ-M)-DARPin were pre-incubated in pH 7.0 or 8.0 of 100 mM Tris-HCl, 150 NaCl or PBS, pH 7.0. Then, different concentrations of Con1 (0.3-6 µM) were mixed with the substrate (30 µM) at a certain pH/buffer and incubated for 4 hours at 25°C. The mixtures were then resolved on 10% SDS-PAGE to evaluate the His tag cleavage.

#### 2.2.7. Temperature dependency of Con1 activity

##### A- FRET assay

We assessed the effect of different temperatures on Con1 activity. The Con1 protease and the FRET substrate were pre-incubated in buffer A at different temperatures (4, 15, 25, 37, 50 and 70°C) for 10 min. The reaction was initiated by mixing Con1 with substrate to final concentrations of 0.5 µM and 2.5 µM, respectively and incubation continued for further 2 hours. The end-point fluorescence was measured at 480 and 520 nm and the amount of cleavage was calculated.

##### B- SDS-PAGE experiment

Con1 and His7-(EAVYHQ-M)-DARPin were pre-incubated of 100 mM Tris-HCl, 150 NaCl, pH 7.0 at different temperatures (0, 4, 12, 25, 37 or 50°C). Then, 1 µM of Con1 was mixed with the 100 µM substrate and incubated for 4 hours at the specified temperature. The mixtures were then resolved on 10% SDS-PAGE to evaluate the His tag cleavage.

#### 2.2.8. His tag removal from different substrates

We used Con1 to cleave an N-terminal His purification tag from different POI. We mixed Con1 (0.5 µM) with the His7 tagged POI (50 µM). Tested POI include His7-(EAVYHQ-M)-StefinA, His7-(EAVYHQ-M)-Interferonα2, His7-(EAVYHQ-G)-DARPin and His7-(EAVYHQ-M)-GFP. The mixtures were incubated at room temperature, overnight with gentle shaking. The cleavage was assessed by separating proteins on SDS-PAGE.

The P1’ tolerance was examined by incubating 0.5 µM of Con1 or Δ222-234Con1 with 50 µM of different His7-(EAVYHQ-X)-DARPin substrates with 12 different N-terminals and incubated overnight at room temperature. The cleavage was assessed by separating proteins on SDS-PAGE.

#### 2.2.9. Investigating different molar ratios and incubation times

- Increasing Con1 concentrations (0.1, 0.2, 1, 2 and 10 µM) were mixed with 100 µM of His_7_-(EAVYHQ-X)-DARPin; where X = Gly, Asp, Met or Ile. The reactions were incubated at 25°C for either 4 hours or 17 hours.
- Increasing His_7_-(EAVYHQ-X)-DARPin; where X = Asp or Met (100, 200, 300, 400, 500 or 600 µM) were mixed with 1 µM of Con1 and the reactions were incubated at 25°C for 17 hours.
- 1 µM of Con1 was mixed with 100 µM of His_7_-(EAVYHQ-X)-DARPin; where X = Gly, Asp, Met or Ile and the reactions were incubated at 25°C and fractions were collected at certain time points (0, 0.5, 1, 2, 4 and 24 hours).

The cleavage was assessed by separating proteins on SDS-PAGE.

## 3. Results and Discussion

### 3.1. Identification of TEV-like proteases

We used a FASTA search to identify TEV-like proteases in the UniProt Knowledgebase and UniProtKB/ Swiss-Prot isoforms databases [14]. From the list of 50 proteases identified, we excluded those with more than 90% or less than 50% identity to TEV protease. Eight proteases meet these criteria ranging in sequence identity with TEV protease from 52.3-60.6% (Table 1). All were plant viral proteases, of the type Nuclear Inclusion protein a (NIa) [13]. In Table 1, we specify the cleavage site, if known, (data taken from the MERPOS database [23]; Supplementary Fig. 1).

**Table 1.** Proteases with ≥ 50% and ≤ 90% identity to TEV protease.

| UniProt ID | Viral origin | Abbreviation | Genome Position | Identity (%) | Cleavage site |
| --- | --- | --- | --- | --- | --- |
| P04517 | *Tobacco etch virus* | TEV | 2038-2279 | Reference | ENLYFQ-S |
| Q85197 | *Potato virus A* | PVA | 2032-2274 | 60.6 | EAVQFQ-S |
| Q02597 | *Turnip mosaic virus (strain Quebec)* | TuMV_Q | 2116-2358 | 54.8 | TVYHQ-A |
| P89509 | *Turnip mosaic virus (strain Japanese)* | TuMV_J | 2117-2359 | 54.8 | EAVYHQ-S |
| Q01681 | *Plum pox potyvirus (strain El amar)* | PPV | 426-668 | 53.7 | TVYHQ-A |
| P20234 | *Ornithogalum mosaic virus* | OMV | 123-365 | 53.3 | -- |
| P09814 | *Tobacco vein mottling virus* | TVMV | 2002-2242 | 52.3 | ETVRFQ-S |
| P89876 | *Lettuce mosaic virus (strain E)* | LMVE | 2215-2457 | 52.9 | -- |
| P31999 | *Lettuce mosaic virus (strain O / French)* | LMV0 | 2215-2457 | 52.5 | -- |

Uniprot ID of the viral polyprotein, name of the virus, the abbreviated name of the virus, location in the viral genome, percentage identity of the viral protease to the TEV protease, optimal substrate cleavage sequence (as reported in MEROPS database), (-) indicate the position of the cleavage between P<sub>1</sub> and P<sub>1</sub>', (--) Not identified.

We assessed the relative solubility of the shortlisted proteases using CamSol analysis of their sequences [15–17]. TuMV-J was predicted to be the most soluble, and PVA the least (Supplementary Table 1). Therefore, the PVA protease was excluded from further analyses.

### 3.2. Consensus protease design

The remaining eight proteases were aligned using Clustal Omega (Fig. 1A) [14, 18], from this alignment, we generated a consensus sequence. The entire consensus sequence is 243 amino acids. In the sequence, 207 residues have more than 40% conservation amongst the eight aligned sequences. For the 36 non-conserved positions, the residues were chosen based on the degree of conservation and hydrophilicity. We created ten sequences (Con1-Con10) with different combinations at certain positions (the 36 non-conserved and additional variable 36 positions) and predicted their solubility using CamSol (Fig. 1B; Supplementary Table 2).

**Figure 1.**
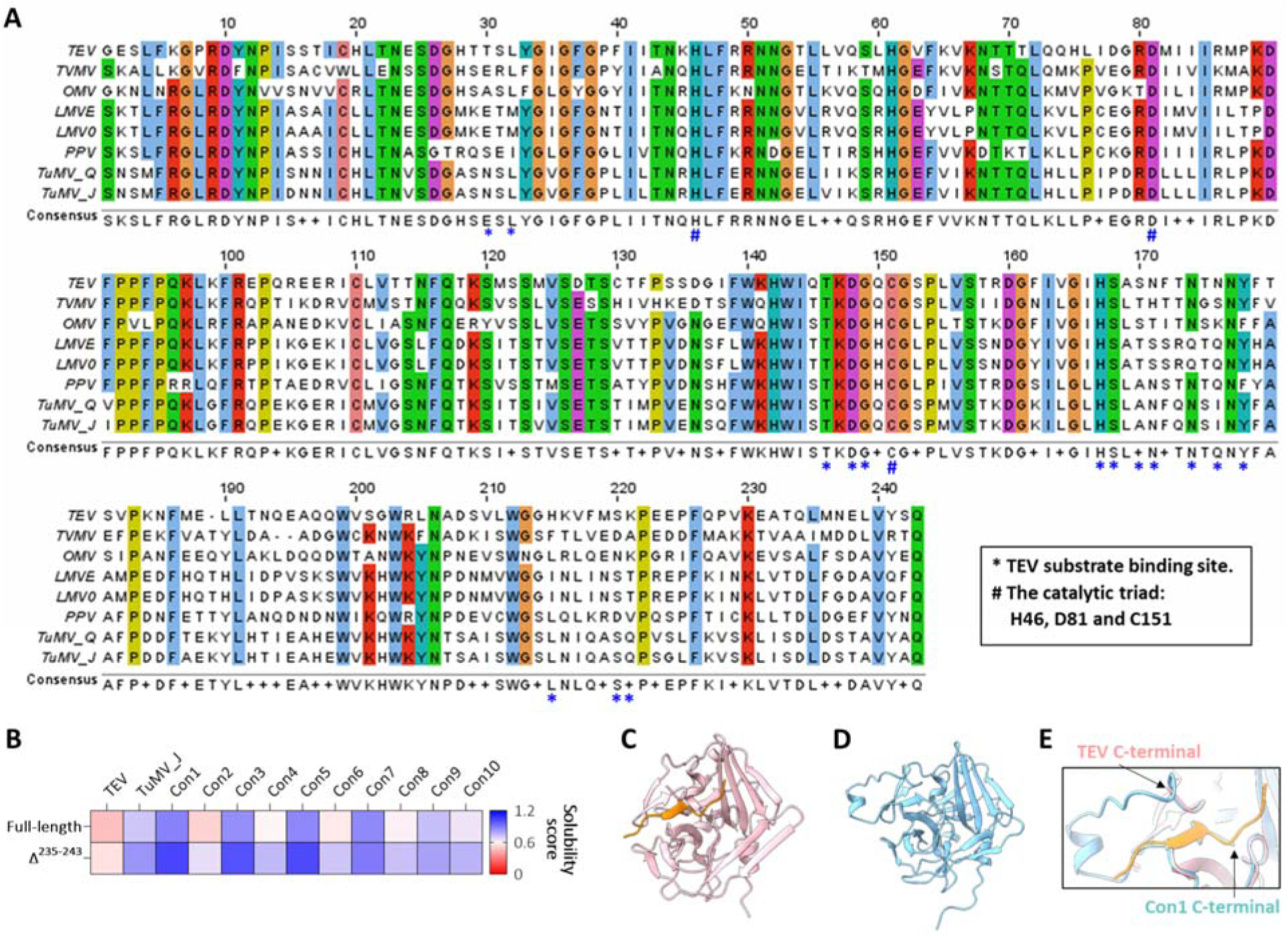
Design of Consensus protease. (**A**) Alignment of the full-length amino acid sequences of the indicated proteases, using Clustal Omega, with a 75% identity threshold coloured by the default Clustal scheme (light blue for hydrophobic, red for +ve charge, magenta for –ve charge, green for polar, pink for Cysteine, orange for Glycine, yellow for Proline, cyan for aromatic and colourless for non-conserved residues). The consensus sequence was automatically defined by the <u>JalView</u> viewer. The asterisk annotations indicate the TEV residues which interact with the substrate residues (P_1_, P_3_, P_6_ and P_1_’). The hashtag annotations indicate the catalytic triad (H46, D81 and C151). (**B**) Intrinsic solubility prediction of the full-length (1-243) and C-terminal truncated (1-234; Δ^235-243^) versions of the designed library along with TEV and TuMV_J proteases as references. The bluer, the more soluble. (**C**) Ribbon representation of TEV protease structure (pink) complexed with uncleaved substrate (orange; PDB ID: 1LVB). (**D**) AlphaFold2 model of the full-length Con1 protease (1-243 aa) predicted by ColabFold. (**E**) Close-up view of the active site of Con1 (light blue) superimposed onto TEV protease (pink) the C-terminus of Con1 is indicated in orange.

We also used Camsol to evaluate the predicted effect of the C-terminal (235-243) deletion on the solubility of Con1. The deletion was predicted to enhance solubility (Fig. 1B). This result is consistent with a form of TEV with the C terminal tail deleted often being used to enhance solubility (Fig. 1C-E) [8, 11, 19].

We designed genes encoding Con1 or TuMV_J with the C-terminal amino acids (235-243 deleted), and codons optimized for expression in E. coli. For TEV protease, the expression plasmid pRK793 (Addgene #8827 [11]) was used. All three proteases were expressed in soluble form and purified as described (Supplementary Fig. 2).

### 3.3. Con1 solubility

A protease with high solubility is a key goal of this study. To experimentally assess protein solubility, we tracked the highest concentration of Con1 or TEV protease that could be achieved by concentrating until the solubility threshold beyond which aggregation occurred [10]. Such experiments indicated that Con1 is at least 2x more soluble than TEV protease: TEV started to aggregate at concentrations above 15 mg/ml whereas Con1 did not aggregate up to 30 mg/ml (Supplementary Fig. 3A). Also, TEV protease aggregated rapidly when high concentration protein samples were incubated at room temperature (Supplementary Fig. 3B).

### 3.4. Con1 cleaves EAVYHQ-S but not ENLYFQ-G

The consensus sites for substrate recognition and cleavage are ENLYFQ-G for TEV (Fig. 2A) and EAVYHQ-S for TuMV_J (Fig. 2B) [9, 19, 24, 25]. We assessed the specificity of TEV, TuMV_J and Con1 using a FRET-based assay in which different potential substrate cleavage sequences were incorporated between cyan fluorescent protein (CFP, donor) and yellow fluorescent protein (YFP, acceptor; Fig. 2C; Supplementary Notes 1.4 and 1.5; Supplementary Fig. 4) [21, 26]. See materials and methods for more details. Con1 and TuMV_J cleave EAVYHQ-S but not ENLYFQ-G (Fig. 2D and E; Supplementary Fig. 5) which is consistent with the reported TuMV_J specificity [24, 27, 28].

**Figure 2.**
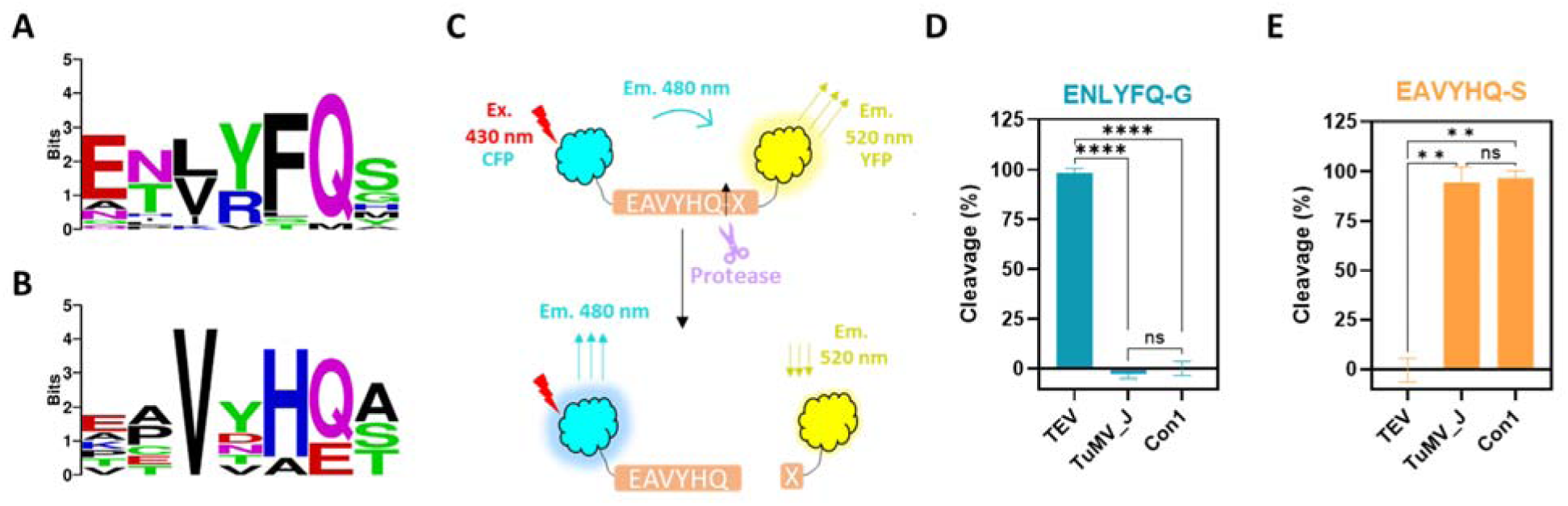
Substrate specificity of Con1 protease. (**A** and **B**) Substrate sequence preferences for cleavage by TEV protease; n=19 (**A**) or TuMV_J protease; n=7 (**B**). The amino acid preferences at each position (P_6_-P_5_-P_4_-P_3_-P_2_-P_1_-P_1_’) are indicated as logo plots (<u>WebLogo</u> online application). (**C**) Schematic illustration describing the FRET-based assay. The substrates contain cyan fluorescent protein (CFP; donor) and yellow fluorescent protein (YFP; acceptor) separated by the potential cleavage site. Before cleavage, by exciting substrate at 430 nm, the emission of YFP is high at 520 nm while CFP emission is 480 nm is low. If the substrate is cleaved by the protease, the YFP emission decreases and CFP emission increases. (**D** and **E**) Screening Con1, TuMV_J or TEV proteases against FRET substrates with different cleavage sites, ENLYFQ-G (**D**) or EAVYHQ-S (**E**). All the results are the mean of 2 technical repeats ± standard deviation. Ordinary one-way ANOVA was used: \*\*\*\**P* < 0.0001; \*\*\**P* < 0.001; \*\**P* < 0.01; \**P* < 0.05; ns, *P* > 0.05 with Tukey’s test correction.

### 3.5. Con1 and TuMV_J cleavage of EAVYHQ-X substrates

We prepared FRET substrates with different potential cleavage sites, EAVYHQ-X (X= alanine, methionine, aspartic or proline). We used the FRET-based assay to determine the kinetic parameters of Con1 and TuMV_J against substrates with different residues at the P1’ position. As expected, neither enzyme cleaves a substrate with proline at the P1’ position (Supplementary Fig. 6). Both Con1 and TuMV_J cleave all the other substrates, with Con1 exhibiting a higher catalytic rate than TuMV_J against them (Fig. 3; Supplementary Fig. 7 and 8). The kinetic parameters of Con1 are indicated in Table 2, but the low catalytic activity of TuMV_J precluded accurate calculations of kinetic parameters. Substrates with different residues at the P1’ position are cleaved at similar rates by Con1, differences are in Km.

**Figure 3.**
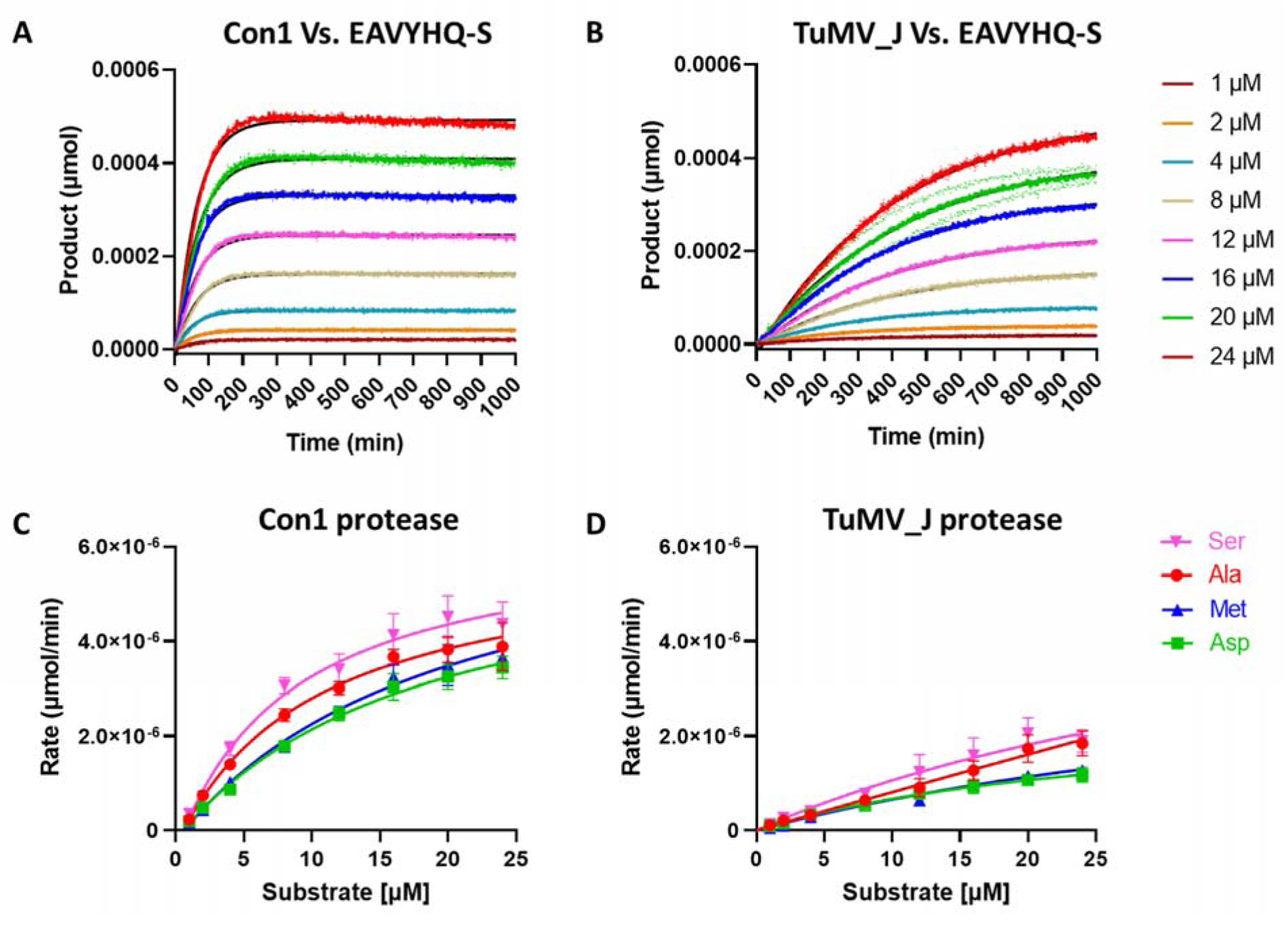
Kinetic studies of Con1 and TuMV_J. (**A** and **B**) Representative plots of the product formation versus time for the indicated concentrations of EAVYHQ-S FRET substrate incubated with 0.4 μM Con1 (**A**) or TuMV_J (**B**). The results show the mean of 2 technical replicates (solid lines) ± standard deviation (dotted lines). The curves were fitted by the exponential plateau model on GraphPad Prism (black lines) to determine the reaction rates. See materials and methods for data analysis and equations. (**C** and **D**) Rate versus substrate concentration plots for Con1 (**C**) and TuMV_J (**D**) against the different FRET substrates (P_1_’=Ser, Ala, Asp or Met). The results are the mean of 2 technical repeats ± standard deviation. The curves are fitted by the Michaelis-Menten non-linear regression using GraphPad Prism software. The curves colors are indicated on (**D**).

**Table 2.**
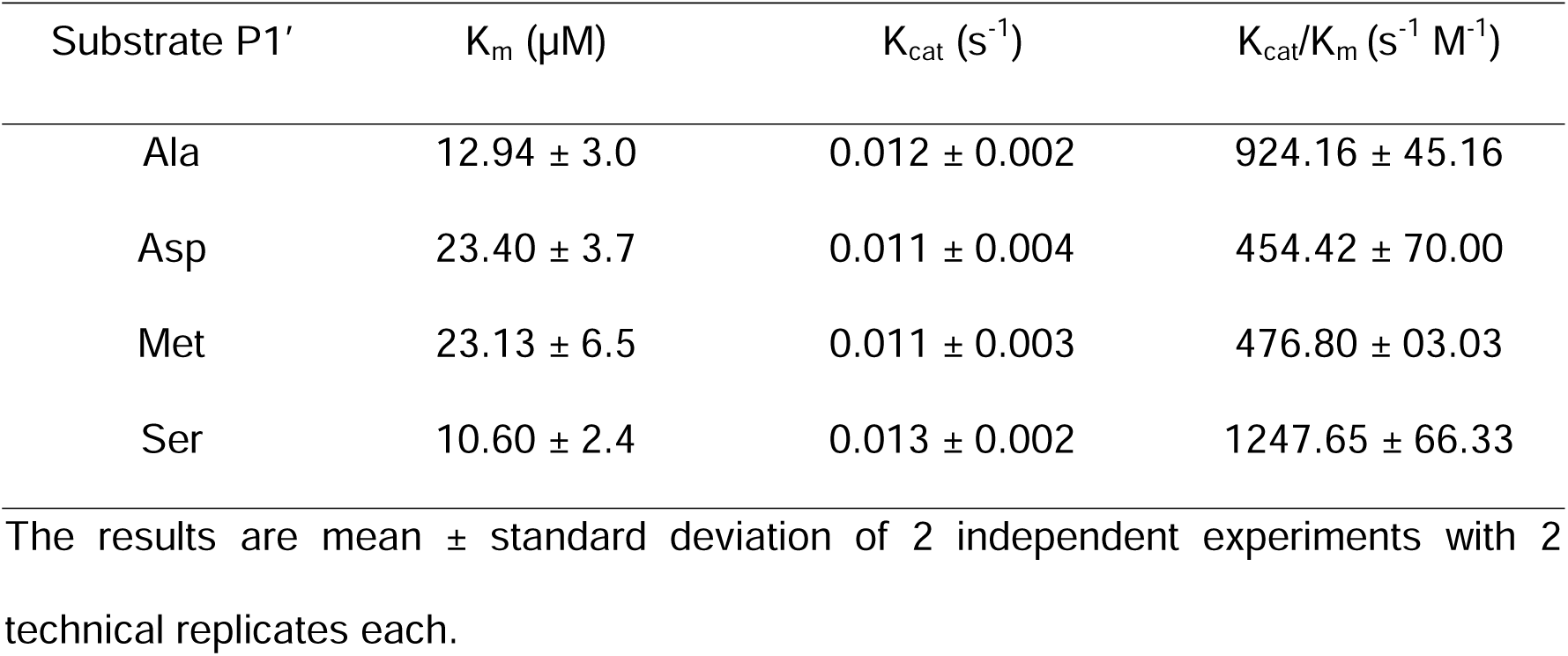
Summary of the kinetic parameters of Con1.

### 3.6. Dependency of Con1 activity on pH and temperature

To ascertain the conditions in which Con1 could be used in practice, we characterized Con1 activity at different pHs, buffers and temperatures. For the pH studies, Con1 protease and substrate were pre-incubated in buffers of the desired pHs including Sodium Citrate (pH 4-6), Tris (pH 7.0-9.0), or Sodium Borate (pH 10), then the reaction was initiated by mixing the substrate with the enzyme. The optimal pH for Con1 activity was determined around pH 7.0 (Fig. 4A; Supplementary Fig. 9A) without any difference observed between Tris-HCl or PBS buffers representing pH 7.0 (Supplementary Fig. 9B).

**Figure 4.**
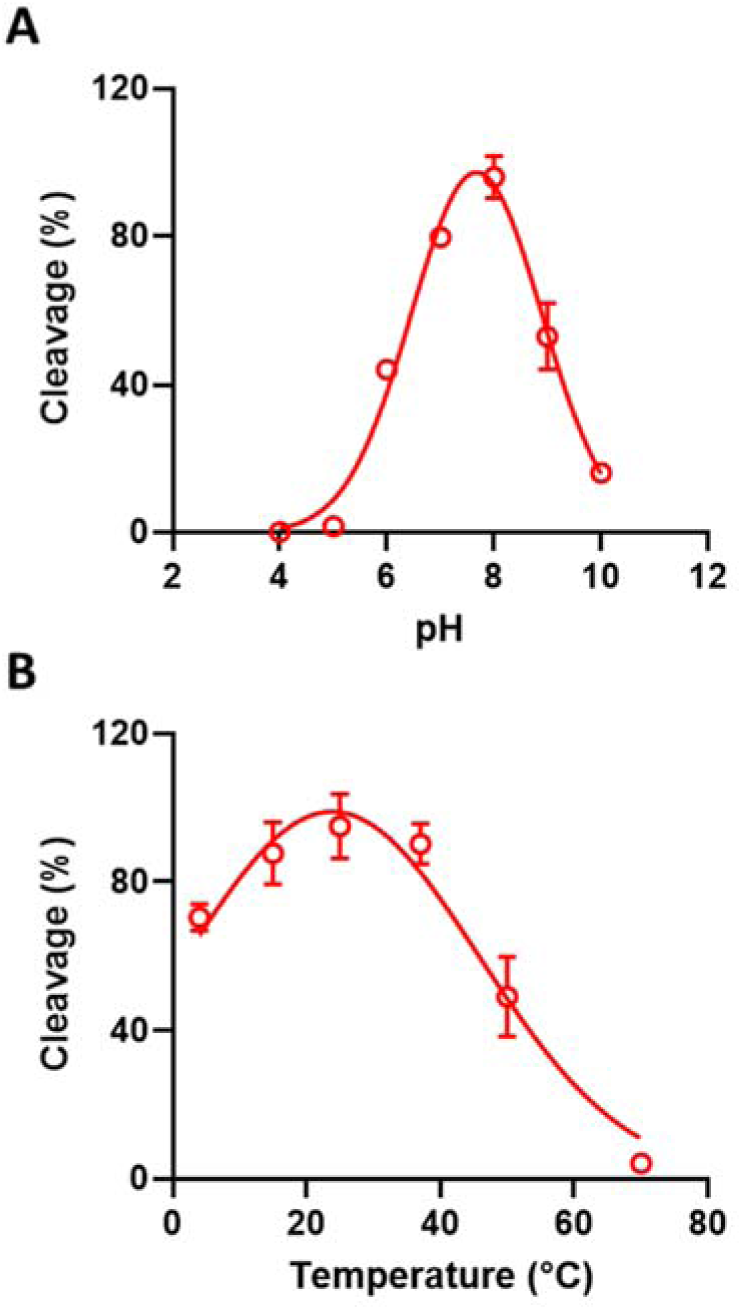
Characterization the pH and temperature dependency of Con1 activity. (**A**) Con1 and FRET substrate (EAVYHQ-S) were mixed to initiate the reaction after 10 min equilibration at the desired pH. The amount of FRET substrate cleaved after 2 hours incubation at room temperature at a given pH was determined. Results are expressed as a percentage of the cleavage that occurred at the optimal pH (pH 7.7). (**B**) Con1 and substrate were equilibrated in buffer A at the specified temperature for 10 min and the reaction was initiated by mixing them. The amount of FRET substrate cleaved after 2 hours incubation was determined. Results are expressed as a percentage of the cleavage that occurred at the optimal temperature (24°C). All the results are the mean of 2 technical repeats ± standard deviation.

We further assessed the kinetics of Con1 at different pH (6.0, 7.0, 8.0 and 9.0) against the FRET substrate with the Met at P1’ position. The best kinetic parameters were observed at pH 7.0 with a lower K_m_ and 2x higher cleavage efficiency than pH 8.0 (Supplementary Fig. 10). Con1 was very slow at pH 6.0 and 9.0 at which the kinetic parameters couldn’t be calculated.

To investigate the effect of temperature on the proteolysis, Con1 and substrate were pre-incubated for 10 minutes at various temperatures (4, 15, 25, 37, 50 or 70°C) in buffer A. Then, the proteolytic reaction was initiated by mixing the substrate with the enzyme. The results indicate that Con1 activity increased by raising the incubation temperature to 37°C beyond which the enzyme started to die, and the optimum activity was determined at a temperature of 24°C (Fig. 4B; Supplementary Fig. 9C).

### 3.7. Con1 protease cleaves His tag independent of the identity of the P1’ residue

A key goal of this study was to develop a protease that could be used to remove N-terminal affinity purification tags from a protein of interest (POI) after affinity purification. Therefore, we designed different His tagged proteins of interest (POI) with P6-P1 of the Con1 recognition site (EAVYHQ) directly upstream of the POI. The first amino acid of the POI contributes to the P1’ residue. These POI are the human cysteine proteinase inhibitor, Stefin A [29], the human cytokine, Interferon α2 [30], a designed ankyrin repeat protein (DARPin) [31] and green fluorescent protein (GFP) [32] (Supplementary Notes 1.10-1.13; Supplementary Fig. 11). After affinity purification using Ni-NTA, the His tagged POI was incubated overnight with Con1. The removal of the N-terminal purification tag was then assessed by separating the proteins using SDS PAGE. As seen in Figure 5A, the tags are all completely removed, and there is no undesired additional cleavage observed.

**Figure 5.**
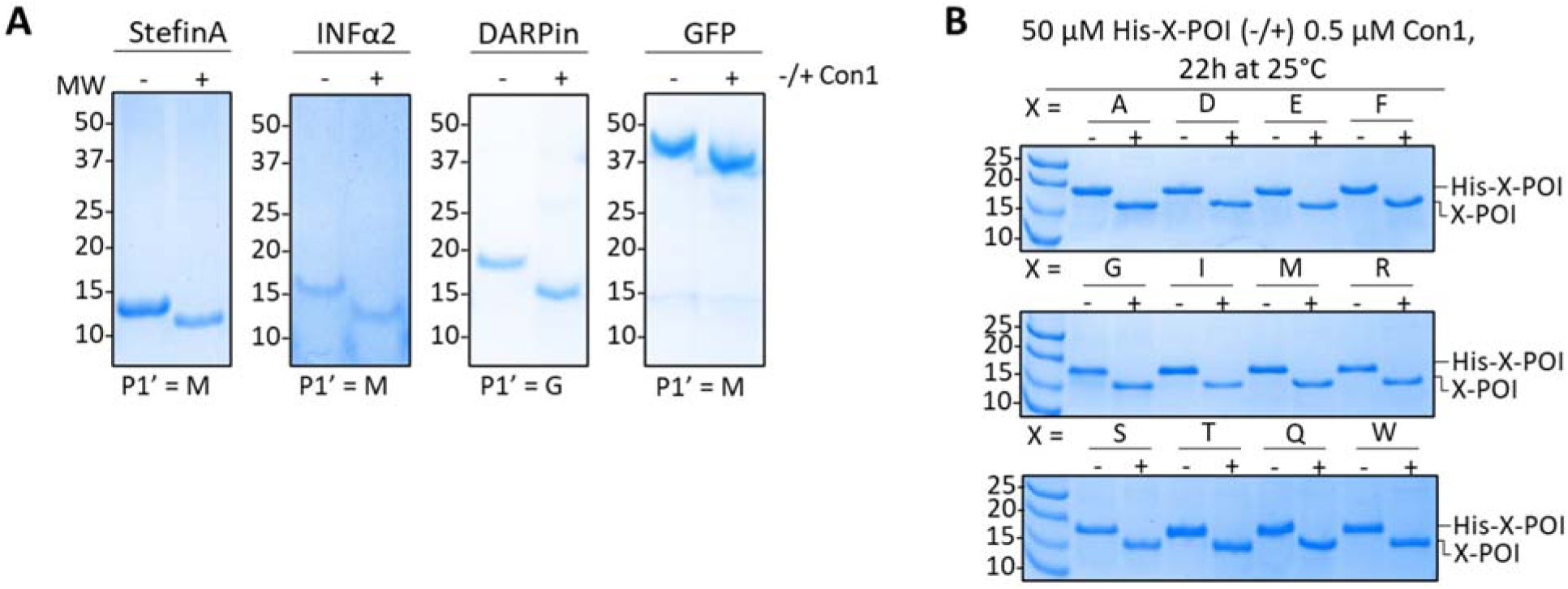
SDS PAGE analysis of different His tagged proteins of interest before (–) and after (+) incubation with protease. (A) Evaluation of His tag removal by Con1 from different substrates as indicated. (B) Screening the P1’ tolerance of Con1 against His tagged DARPins proteins with different N-terminal as indicated. Complete removal of the purification tag is evident by the decrease in MW of the protein. The reaction mixtures contained 0.5 µM Con1 and 50 µM substrate final concentrations and were incubated overnight at room temperature.

To screen the P1’ tolerance of Con1, we mutated the natural N-terminal amino acid of the DARPin (Gly) to 11 different residues covering different grouping of amino acids: hydrophobic amino acids (A, F, I, M and W), polar uncharged amino acids (Q, S and T), positively charged amino acids (R) and negatively charged amino acids (D and E; Supplementary Fig. 12). Con1 efficiently cleaved the His tag from all the studied substrates with different N-termini (acting as a P1’ position). This result indicates that Con1 can cleave off N-terminal tags, independent of the P1’ position (i.e., the N-terminal amino acid of the POI) (Fig. 5B).

Additional experiments were performed to evaluate the effects of changing the enzyme to substrate molar ratio (1: 1000, 1: 600, 1: 500, 1: 400, 1: 300, 1: 200, 1: 100, 1: 50 and 1: 10) versus the incubation time (4h and 17 h) to achieve 100% cleavage of a tag from selected P1’ residues (Gly, Asp, Met or Ile). We found that the His tag was fully cleaved from the Gly and Asp substrates after 2 hours of incubation at a 1: 100 molar ratio, a longer incubation at the same ratio resulted in the cleavage of the tag from the Met and Ile substrates (Supplementary Fig. 13).

### 3.8. Con1 activity is C-terminal sensitive

Con1 is 234 amino acids long. The predicted AlphaFold2 model of the structure indicated the C-terminal residues 222-234 to be unstructured (Supplementary Fig. 14A). This motivated us to study the effect of C-terminal deletion on the activity of the Con1. We prepared two protease versions by deleting 228-234 (^Δ228-234^Con1) or 222-234 (^Δ222-234^Con1) residues (Supplementary Fig. 14B). Kinetic studies of Con1, ^Δ228-234^Con1 and ^Δ222-234^Con1 against FRET substrate with the EAVYHQ-M cleavage site shows better cleavage by ^Δ222-234^Con1 than by the original Con1. Cleavage by ^Δ228-234^Con1 is worse than cleavage by ^Δ222-234^Con1 (Fig. 6A; Supplementary Fig. 15). The superior catalytic activity of ^Δ222-234^Con1 derives from a K_m_ value that is about half that of the original Con1. Both have similar k_cat_ values (Table 3). We speculate that because the C-terminus is close to the active site, the unstructured tail may interfere in some fashion with substrate binding.

**Figure 6.**
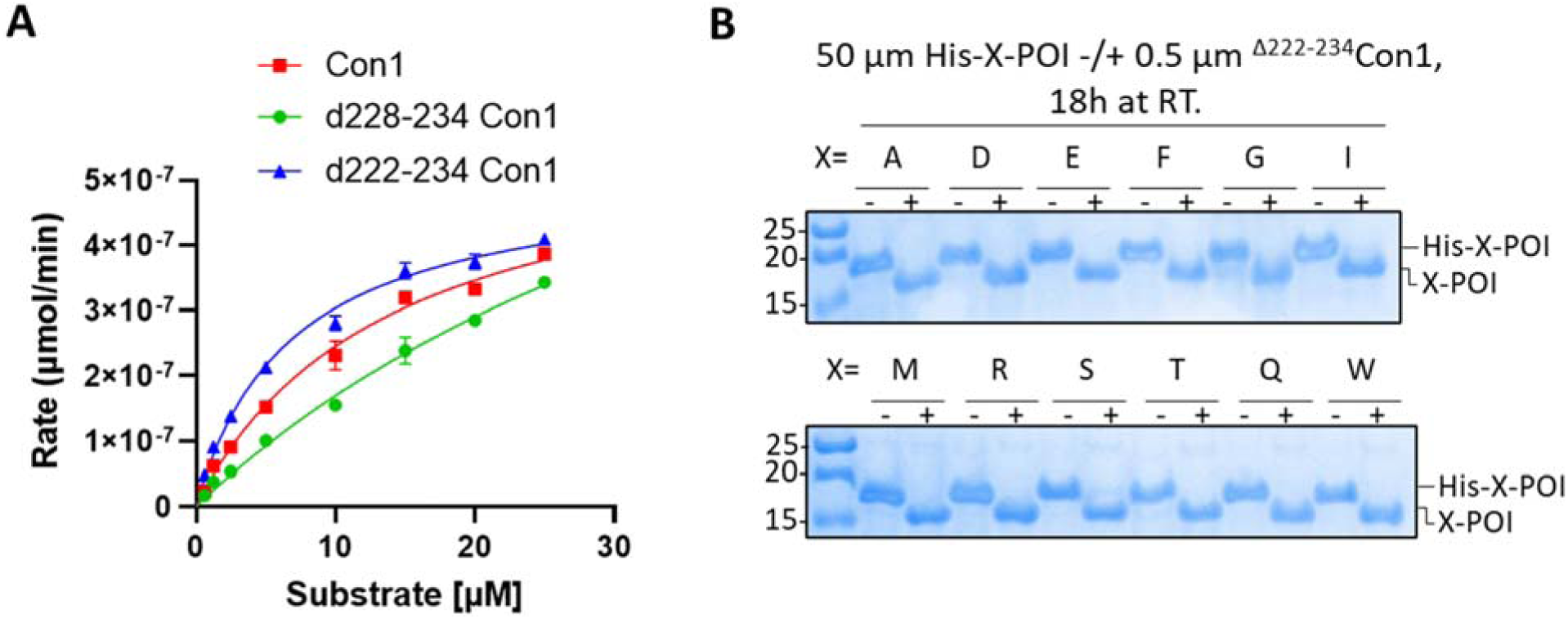
Kinetic studies of the C-terminal truncated versions of Con1. (A) Kinetics summary of Con1, ^Δ228-234^Con1 and ^Δ222-234^Con1 against EAVYHQ-M FRET substrate. The results are the mean of 2 technical repeats ± standard deviation. The curves are fitted by the Michaelis-Menten non-linear regression using GraphPad Prism software. (B) Screening the P1’ tolerance of ^Δ222-234^Con1 against His tagged DARPins proteins with different N-terminal as indicated. Complete removal of the purification tag is evident by the decrease in MW of the protein. The reaction mixture contained 0.5 µM Con1 and 50 µM substrate final concentrations and was incubated overnight at room temperature.

**Table 3.**
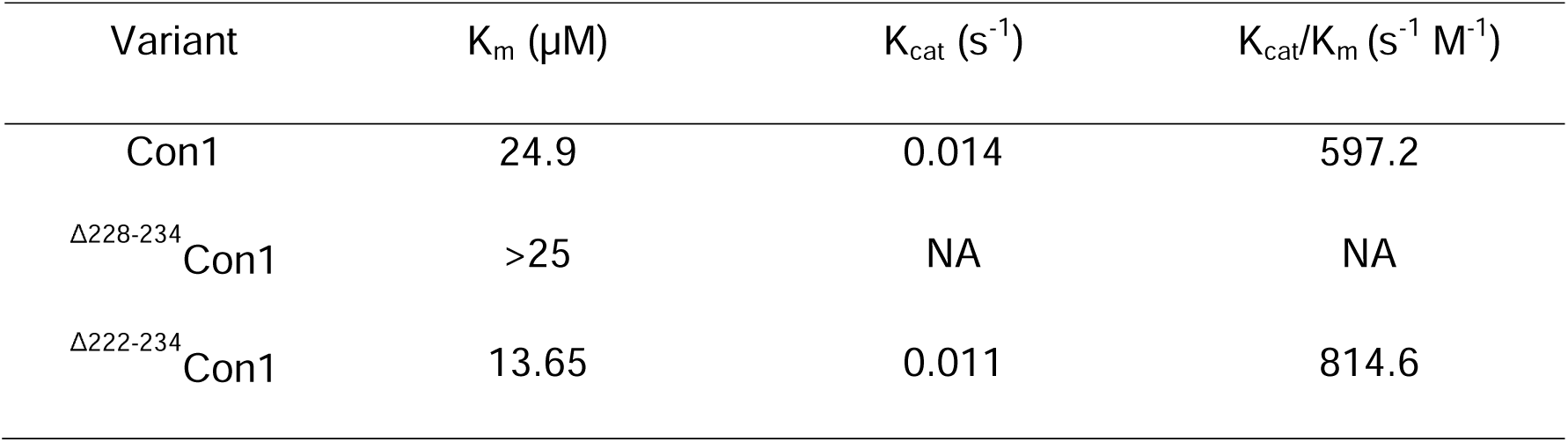
Summary of the kinetic parameters of truncated versions of Con1.

Most importantly ^Δ222-234^Con1 exhibits the same P1’ tolerance as Con1. We screened ^Δ222-234^Con1 activity against the 12 different DARPins with the different N-termini (Supplementary Fig. 12). ^Δ222-234^Con1 efficiently cleaved the His tag from all the substrates (Fig. 6B).

## 4. Conclusion

Affinity purification of proteins is a powerful method, applicable to a wide range of POI. For many applications, it is necessary to remove the affinity tag after purification. TEV protease was one of the first exploited for this purpose, and it is widely used. Two features of TEV protease that are less than ideal, are its decreased activity against cleavage sequences with residues other than Gly or Ser in the P_1_’ position, and its low solubility. Inspired by TEV protease, we sought to create a protease that has activity against substrates with different residues at the P_1_’ position, and which is significantly more soluble than TEV protease. Here we have described the properties of new enzymes, Con1 proteases, whose substrate specificity and solubility make them well-suited for the removal of N-terminal affinity purification tags from a wide variety of proteins.

## CRediT authorship contribution statement

**RS** and **LR** led and designed the study. **RS** performed the computational and experimental work, analysed data, generated figures, and wrote the first draft of the manuscript. **LR** conceptualized the study, analysed data and edited the manuscript.

## Data availability

Data will be made available on request.

## Acknowledgement

This work was funded by an Industrial Biotechnology Innovation Centre (IBioIC) Innovator Award and the BBSRC Impact Acceleration Account (IAA) fund (BBSRC IAA PIII117) in partnership with Fujifilm Diosynth Biotechnology UK (FDBK). The University of Edinburgh is a member of the Centre of Excellence in Bioprocessing, along with Fujifilm Diosynth Biotechnology (FDB), the University of York and the University of Manchester. We were motivated to perform this work by Dr Ray O’Donnell (FDBK). We acknowledge the importance of frequent interactions with Dr Christopher Lennon, Dr Kenneth Holbourn and Dr Nicola Preston (FDBK). We thank Dr Mai-Brit Jensen, Dr Ella Thornton and Kasia Stefaniak (University of Edinburgh) for their thoughtful comments on the manuscript.

## References

[1] D.S. Waugh, Making the most of affinity tags, Trends Biotechnol 23(6) (2005) 316–20. 10.1016/j.tibtech.2005.03.012.

[2] S. Raran-Kurussi, D.S. Waugh, The ability to enhance the solubility of its fusion partners is an intrinsic property of maltose-binding protein but their folding is either spontaneous or chaperone-mediated, PLoS One 7(11) (2012) e49589. 10.1371/journal.pone.0049589.

[3] F. Schafer, N. Seip, B. Maertens, H. Block, J. Kubicek, Purification of GST-Tagged Proteins, Methods Enzymol 559 (2015) 127–39. 10.1016/bs.mie.2014.11.005.

[4] S. Costa, A. Almeida, A. Castro, L. Domingues, Fusion tags for protein solubility, purification and immunogenicity in Escherichia coli: the novel Fh8 system, Front Microbiol 5 (2014) 63. 10.3389/fmicb.2014.00063.

[5] N.T.P. Le, T.T.P. Phan, H.T.T. Phan, T.T.T. Truong, W. Schumann, H.D. Nguyen, Influence of N-terminal His-tags on the production of recombinant proteins in the cytoplasm of Bacillus subtilis, Biotechnol Rep (Amst) 35 (2022) e00754. 10.1016/j.btre.2022.e00754.

[6] H. Block, B. Maertens, A. Spriestersbach, J. Kubicek, F. Schafer, Proteolytic Affinity Tag Cleavage, Methods Enzymol 559 (2015) 71–97. 10.1016/bs.mie.2014.11.009.

[7] D.S. Waugh, An overview of enzymatic reagents for the removal of affinity tags, Protein Expr Purif 80(2) (2011) 283–93. 10.1016/j.pep.2011.08.005.

[8] S. Raran-Kurussi, S. Cherry, D. Zhang, D.S. Waugh, Removal of Affinity Tags with TEV Protease, Methods Mol Biol 1586 (2017) 221–230. 10.1007/978-1-4939-6887-9_14.

[9] R.B. Kapust, J. Tozser, T.D. Copeland, D.S. Waugh, The P1’ specificity of tobacco etch virus protease, Biochem Biophys Res Commun 294(5) (2002) 949–55. 10.1016/S0006-291X(02)00574-0.

[10] L.D. Cabrita, D. Gilis, A.L. Robertson, Y. Dehouck, M. Rooman, S.P. Bottomley, Enhancing the stability and solubility of TEV protease using in silico design, Protein Sci 16(11) (2007) 2360–7. 10.1110/ps.072822507.

[11] R.B. Kapust, J. Tozser, J.D. Fox, D.E. Anderson, S. Cherry, T.D. Copeland, D.S. Waugh, Tobacco etch virus protease: mechanism of autolysis and rational design of stable mutants with wild-type catalytic proficiency, Protein Eng 14(12) (2001) 993–1000. 10.1093/protein/14.12.993.

[12] J.E. Tropea, S. Cherry, D.S. Waugh, Expression and purification of soluble His(6)-tagged TEV protease, Methods Mol Biol 498 (2009) 297–307. 10.1007/978-1-59745-196-3_19.

[13] C. UniProt, UniProt: the Universal Protein Knowledgebase in 2023, Nucleic Acids Res 51(D1) (2023) D523–D531. 10.1093/nar/gkac1052.

[14] F. Madeira, M. Pearce, A.R.N. Tivey, P. Basutkar, J. Lee, O. Edbali, N. Madhusoodanan, A. Kolesnikov, R. Lopez, Search and sequence analysis tools services from EMBL-EBI in 2022, Nucleic Acids Res 50(W1) (2022) W276–9. 10.1093/nar/gkac240.

[15] P. Sormanni, M. Vendruscolo, Protein Solubility Predictions Using the CamSol Method in the Study of Protein Homeostasis, Cold Spring Harb Perspect Biol 11(12) (2019). 10.1101/cshperspect.a033845.

[16] P. Sormanni, F.A. Aprile, M. Vendruscolo, The CamSol method of rational design of protein mutants with enhanced solubility, J Mol Biol 427(2) (2015) 478–90. 10.1016/j.jmb.2014.09.026.

[17] P. Sormanni, L. Amery, S. Ekizoglou, M. Vendruscolo, B. Popovic, Rapid and accurate in silico solubility screening of a monoclonal antibody library, Sci Rep 7(1) (2017) 8200. 10.1038/s41598-017-07800-w.

18. A.M. Waterhouse, J.B. Procter, D.M. Martin, M. Clamp, G.J. Barton, Jalview Version 2--a multiple sequence alignment editor and analysis workbench, Bioinformatics 25(9) (2009) 1189–91. 10.1093/bioinformatics/btp033.

[19] J. Phan, A. Zdanov, A.G. Evdokimov, J.E. Tropea, H.K. Peters, 3rd, R.B. Kapust, M. Li, A. Wlodawer, D.S. Waugh, Structural basis for the substrate specificity of tobacco etch virus protease, J Biol Chem 277(52) (2002) 50564–72. 10.1074/jbc.M207224200.

[20] H.M. Beyer, P. Gonschorek, S.L. Samodelov, M. Meier, W. Weber, M.D. Zurbriggen, AQUA Cloning: A Versatile and Simple Enzyme-Free Cloning Approach, PLoS One 10(9) (2015) e0137652. 10.1371/journal.pone.0137652.

[21] M.H.T.R.T. Hay, FRET-Based In Vitro Assays for the Analysis of SUMO Protease Activities, in: H.D. Ulrich (Ed.), METHODS IN MOLECULAR BIOLOGY: SUMO Protocols, Humana Press, Totowa, NJ., 2009, pp. 253–268.

[22] D.H. Kim, D.C. Hwang, B.H. Kang, J. Lew, K.Y. Choi, Characterization of Nla protease from turnip mosaic potyvirus exhibiting a low-temperature optimum catalytic activity, Virology 221(1) (1996) 245–9. 10.1006/viro.1996.0372.

[23] N.D. Rawlings, A.J. Barrett, P.D. Thomas, X. Huang, A. Bateman, R.D. Finn, The MEROPS database of proteolytic enzymes, their substrates and inhibitors in 2017 and a comparison with peptidases in the PANTHER database, Nucleic Acids Res 46(D1) (2018) D624–D632. 10.1093/nar/gkx1134.

[24] H. Kang, Y.J. Lee, J.H. Goo, W.J. Park, Determination of the substrate specificity of turnip mosaic virus NIa protease using a genetic method, J Gen Virol 82(Pt 12) (2001) 3115–3117. 10.1099/0022-1317-82-12-3115.

[25] D.-H.K. Ji Seon Han, Kwan Yong Choi, Potyvirus NIa Protease, in: N.R. Alan Barrett, J. Woessner (Ed.), Handbook of Proteolytic Enzymes, Elsevier2012, pp. 2427–2432.

[26] H. Gu, S. Lalonde, S. Okumoto, L.L. Looger, A.M. Scharff-Poulsen, A.R. Grossman, J. Kossmann, I. Jakobsen, W.B. Frommer, A novel analytical method for in vivo phosphate tracking, FEBS Lett 580(25) (2006) 5885–93. 10.1016/j.febslet.2006.09.048.

[27] D.H. Kim, Y.S. Park, S.S. Kim, J. Lew, H.G. Nam, K.Y. Choi, Expression, purification, and identification of a novel self-cleavage site of the Nla C-terminal 27-kDa protease of turnip mosaic potyvirus C5, Virology 213(2) (1995) 517–25. 10.1006/viro.1995.0024.

[28] H.E. Han, S. Sellamuthu, B.H. Shin, Y.J. Lee, S. Song, J.S. Seo, I.S. Baek, J. Bae, H. Kim, Y.J. Yoo, Y.K. Jung, W.K. Song, P.L. Han, W.J. Park, The nuclear inclusion a (NIa) protease of turnip mosaic virus (TuMV) cleaves amyloid-beta, PLoS One 5(12) (2010) e15645. 10.1371/journal.pone.0015645.

[29] J.R. Martin, R. Jerala, L. Kroon-Zitko, E. Zerovnik, V. Turk, J.P. Waltho, Structural characterisation of human stefin A in solution and implications for binding to cysteine proteinases, Eur J Biochem 225(3) (1994) 1181–94. 10.1111/j.1432-1033.1994.1181b.x.

[30] C. Thomas, I. Moraga, D. Levin, P.O. Krutzik, Y. Podoplelova, A. Trejo, C. Lee, G. Yarden, S.E. Vleck, J.S. Glenn, G.P. Nolan, J. Piehler, G. Schreiber, K.C. Garcia, Structural Linkage between Ligand Discrimination and Receptor Activation by Type I Interferons, Cell 146(4) (2011) 621–632. 10.1016/j.cell.2011.06.048.

[31] M.T. Stumpp, H.K. Binz, P. Amstutz, DARPins: a new generation of protein therapeutics, Drug Discov Today 13(15-16) (2008) 695–701. 10.1016/j.drudis.2008.04.013.

[32] S.J. Remington, Green fluorescent protein: a perspective, Protein Sci 20(9) (2011) 1509–19. 10.1002/pro.684.

