## Supplementary materials for "A new highly specific and soluble protease for precise removal of N-terminal purification tags"

**Content:**

1- Supplementary notes (Gene’s sequences):
   1.1. Con1 synthetic gene.
   1.2. TuMV_J synthetic gene.
   1.3. TEV clone (pRK793; Addgene #8827).
   1.4. FRET substrate with EAVYHQ-S site.
   1.5. ENLYFQ-G cleavage site.
   1.6. EAVYHQ-A cleavage site.
   1.7. EAVYHQ-D cleavage site.
   1.8. EAVYHQ-M cleavage site.
   1.9. EAVYHQ-P cleavage site.
   1.10. Stefin A sequence.
   1.11. Interferon α2 sequence.
   1.12. DARPin sequence.
   1.13. Green Fluorescent Protein (GFP)-TRAP sequence.
2- Supplementary Tables:
2.1. Supplementary Table 1. Oligonucleotides.
2.2. Supplementary Table 2. Predicted solubility profiles of the selected proteases.
2.3. Supplementary Table 3. Amino acid sequences of the designed Con1-Con10.
3- Supplementary Figures:
3.1. Supplementary Figure 1. Conserved cleavage sites of proteases.
3.2. Supplementary Figure 2. Purification of Con1, TuMV_J and TEV proteases.
3.3. Supplementary Figure 3. Solubility profiling of Con1 Vs. TEV proteases.
3.4. Supplementary Figure 4. Purification of the FRET substrates.
3.5. Supplementary Figure 5. Screening substrate specificity of Con1, TuMV_J and TEV.
3.6. Supplementary Figure 6. Screening specificity of Con1 and TuMV_J against EAVYHQ-P.
3.7. Supplementary Figure 7. Kinetic studies of Con1 against different P_1_’ substrates.
3.8. Supplementary Figure 8. Kinetic studies of TuMV_J against different P_1_’ substrates.
3.9. Supplementary Figure 9. Characterization of the pH and temperature dependency of Con1 activity.
3.10. Supplementary Figure 10. Kinetic studies of Con1 against FRET-M at different pH.
3.11. Supplementary Figure 11. Purification of His tagged protein of interest (POI).
3.12. Supplementary Figure 12. Purification of His tagged DARPins with different N-terminal.
3.13. Supplementary Figure 13. SDS-PAGE experiments showing the His tag removal by Con1 at different conditions.
3.14. Supplementary Figure 14. Purification of C-terminal truncated versions of Con1.
3.15. Supplementary Figure 15. Kinetic studies of different Con1 versions against FRET-M.

**1- Supplementary notes (Gene sequences):**

1.1. **Con1 (726 bp; IDT, Belgium): (7xHis-Con1)**

atgcatcatcatcatcatcatcacAGCAAATCTTTATTCCGTGGATTACGCGATTACAACCCGATCGCCAGCAATATTTGTCATTTGACTAATGAGAGCGACGGTCATAGTAACTCGTTGTATGGCATCGGGTTTGGGCCGCTTATTATTACAAACCAACATTTGTTTCGTCGCAACAACGGGGAACTTACGATTCAATCTCGCCATGGCGAGTTCGTAGTTAAGAACACCACGCAACTTAAGCTTCTGCCAATCGATGGGCGTGATATCCTTATCATTCGTTTGCCCAAAGATTTCCCACCCTTTCCACAGAAACTTAAATTCCGCCAACCTGAAAAAGGCGAACGCATTTGCTTGGTTGGCTCGAATTTTCAGACCAAGTCCATCACCAGTACAGTCAGCGAAACATCAACAACCATGCCGGTGGAGAATAGTCAATTTTGGAAGCACTGGATTTCAACCAAAGACGGTCATTGTGGCCTGCCTCTTGTTTCCACGAAGGATGGCAAGATCCTGGGCATCCATTCTTTGGCGAATTTTACCAATACGATCAACTACTTCGCCGCATTCCCAGAAGATTTTGAGGAGACGTATCTTCATACCCAAGAAGCTCAAGAATGGGTAAAGCACTGGAAATATAACCCGGATGCGATCTCCTGGGGGAGCCTTAACCTTCAAGAAAGCCAACCAGAAGAGCCTTTCAAAATCGTAAAATTAGTAACGGAC

1.2. **TuMV_J (732 bp; IDT, Belgium): (7xHis-TuMV_J)**

atgcatcatcatcatcatcatcacTCGAATTCTATGTTCCGTGGGTTACGCGATTACAACCCGATTGCCAATAATATTTGCCATTTAACGAACGTGTCCGATGGCGCATCGAACTCTCTGTATGGAGTCGGATTTGGGCCGTTAATCTTAACCAATCGTCATCTGTTCGAACGCAATAATGGGGAATTGGTAATTAAGTCTCGTCACGGTGAATTTGTCATCAAGAATACTACCCAGTTGCACCTTTTACCGATTCCCGATCGTGACTTGCTGCTGATTCGCCTGCCCAAGGACATCCCCCCGTTCCCACAAAAACTGGGGTTCCGCCAACCCGAAAAGGGGGAGCGCATTTGTATGGTTGGCTCCAATTTTCAGACAAAGAGTATCACAAGTGTAGTCTCGGAAACGAGCACAATCATGCCCGTCGAGAACAGTCAGTTTTGGAAGCACTGGATTAGCACGAAGGATGGGCAATGCGGTCTGCCTATGGTTAGCACCAAGGACGGTAAAATTCTGGGGTTACATAGTTTAGCCAATTTTCAAAACTCCATCAATTATTTCGCCGCCTTTCCAGACGACTTTGCTGAAAAATATCTGCATACGATCGAGGCACACGAGTGGGTCAAGCATTGGAAATACAACACATCTGCAATTTCATGGGGATCGCTGAACATTCAAGCATCACAACCGGGTGGCTTATTTAAGGTCAGTAAACTTATTTCCGACTTAGAT

1.3. **TEV (747 bp): (7xHis-TEV)**

ggtcatcatcatcatcatcatcatGGAGAAAGCTTGTTTAAGGGGCCGCGTGATTACAACCCGATATCGAGCACCATTTGTCATTTGACGAATGAATCTGATGGGCACACAACATCGTTGTATGGTATTGGATTTGGTCCCTTCATCATTACAAACAAGCACTTGTTTAGAAGAAATAATGGAACACTGTTGGTCCAATCACTACATGGTGTATTCAAGGTCAAGAACACCACGACTTTGCAACAACACCTCATTGATGGGAGGGACATGATAATTATTCGCATGCCTAAGGATTTCCCACCATTTCCTCAAAAGCTGAAATTTAGAGAGCCACAAAGGGAAGAGCGCATATGTCTTGTGACAACCAACTTCCAAACTAAGAGCATGTCTAGCATGGTGTCAGACACTAGTTGCACATTCCCTTCATCTGATGGCATATTCTGGAAGCATTGGATTCAAACCAAGGATGGGCAGTGTGGCAGTCCATTAGTATCAACTAGAGATGGGTTCATTGTTGGTATACACTCAGCATCGAATTTCACCAACACAAACAATTATTTCACAAGCGTGCCGAAAAACTTCATGGAATTGTTGACAAATCAGGAGGCGCAGCAGTGGGTTAGTGGTTGGCGATTAAATGCTGACTCAGTATTGTGGGGGGGCCATAAAGTTTTCATGGTGAAACCTGAAGAGCCTTTTCAGCCAGTTAAGGAAGCGACTCAACTCATGAATCGTCGTCGCCGTCGC

1.4. **FRET substrate with EAVYHQ-S site (1593 bp):**

**(6xHis-linker-CFP-linker-cleavage site-linker-YFP)**

atgcggggttctcatcatcatcatcatcatGGTATGGCTAGCATGACTGGTGGACAGCAAATGGGTCGGGATCTGTACGACGATGACGATAAGGATCCGGGCCGCATGGTGAGCAAGGGCGAGGAGCTGTTCACCGGGGTGGTGCCCATCCTGGTCGAGCTGGACGGCGACGTAAACGGCCACAAGTTCAGCGTGTCCGGCGAGGGCGAGGGCGATGCCACCTACGGCAAGCTGACCCTGAAGTTCATCTGCACCACCGGCAAGCTGCCCGTGCCCTGGCCCACCCTCGTGACCACCCTGACCTGGGGCGTGCAGTGCTTCAGCCGCTACCCCGACCACATGAAGCAGCACGACTTCTTCAAGTCCGCCATGCCCGAAGGCTACGTCCAGGAGCGCACCATCTTCTTCAAGGACGACGGCAACTACAAGACCCGCGCCGAGGTGAAGTTCGAGGGCGACACCCTGGTGAACCGCATCGAGCTGAAGGGCATCGACTTCAAGGAGGACGGCAACATCCTGGGGCACAAGCTGGAGTACAACTACATCAGCCACAACGTCTATATCACCGCCGACAAGCAGAAGAACGGCATCAAGGCCAACTTCAAGATCCGCCACAACATCGAGGACGGCAGCGTGCAGCTCGCCGACCACTACCAGCAGAACACCCCCATCGGCGACGGCCCCGTGCTGCTGCCCGACAACCACTACCTGAGCACCCAGTCCGCCCTGAGCAAAGACCCCAACGAGAAGCGCGATCACATGGTCCTGCTGGAGTTCGTGACCGCCGCCGGGATCGGTACCGTAGGATTTCTAACAGCGACCGAAGCAGTGTATCATCAATCCCTGGATTCCATCACCGTTAACGGTACCGGTGGAATGGTGAGCAAGGGCGAGGAGCTGTTCACCGGGGTGGTGCCCATCCTGGTCGAGCTGGACGGCGACGTAAACGGCCACAAGTTCAGCGTGTCCGGCGAGGGCGAGGGCGATGCCACCTACGGCAAGCTGACCCTGAAGTTCATCTGCACCACCGGCAAGCTGCCCGTGCCCTGGCCCACCCTCGTGACCACCTTCGGCTACGGCCTGCAGTGCTTCGCCCGCTACCCCGACCACATGAAGCAGCACGACTTCTTCAAGTCCGCCATGCCCGAAGGCTACGTCCAGGAGCGCACCATCTTCTTCAAGGACGACGGCAACTACAAGACCCGCGCCGAGGTGAAGTTCGAGGGCGACACCCTGGTGAACCGCATCGAGCTGAAGGGCATCGACTTCAAGGAGGACGGCAACATCCTGGGGCACAAGCTGGAGTACAACTACAACAGCCACAACGTCTATATCATGGCCGACAAGCAGAAGAACGGCATCAAGGTGAACTTCAAGATCCGCCACAACATCGAGGACGGCAGCGTGCAGCTCGCCGACCACTACCAGCAGAACACCCCCATCGGCGACGGCCCCGTGCTGCTGCCCGACAACCACTACCTGAGCTACCAGTCCGCCCTGAGCAAAGACCCCAACGAGAAGCGCGATCACATGGTCCTGCTGGAGTTCGTGACCGCCGCCGGGATCACTCTCGGCATGGACGAGCTGTACAAG

1.5. **ENLYFQ-G site:**

GAAAATCTTTATTTTCAAGGT

1.6. **EAVYHQ-A site:**

GAAGCAGTGTATCATCAAGCG

1.7. **EAVYHQ-D site:**

GAAGCAGTGTATCATCAAGAT

1.8. **EAVYHQ-M site:**

GAAGCAGTGTATCATCAAATG

1.9. **EAVYHQ-P site:**

GAAGCAGTGTATCATCAACCG

1.10. **Stefin A (345 bp): (7xHis-linker-cleavage site-Stefin A)**

atgcatcatcatcatcatcatcacGGCGGTAGCGAAGCAGTGTATCATCAAATGATTCCCGGCGGTTTATCAGAGGCAAAACCCGCGACGCCAGAGATTCAAGAAATCGTCGACAAAGTTAAACCGCAATTGGAAGAGAAGACCAATGAAACTTATGGCAAGTTAGAGGCCGTTCAATACAAGACCCAAGTTGTTGCCGGCACGAATTATTACATTAAGGTTCGTGCGGGCGACAATAAGTACATGCACCTCAAGGTCTTTAAATCATTGCCTGGTCAGAATGAGGATCTCGTTTTAACGGGCTACCAAGTTGACAAGAATAAAGACGACGAACTCACTGGATTT

1.11. **Interferon α2 (549 bp): (7xHis-linker-cleavage site-Interferon α2)**

atgcatcatcatcatcatcatcacGGCGGTAGCGAAGCAGTGTATCATCAAATGTGCGACTTGCCCCAGACACACAGCTTAGGATCTCGCCGTACCCTGATGTTATTAGCGCAAATGCGTCGCATTAGCCTCTTCTCATGCCTCAAGGACCGCCATGACTTTGGGTTCCCACAAGAGGAGTTTGGCAACCAATTCCAGAAGGCAGAGACCATTCCTGTGTTGCACGAGATGATCCAGCAAATTTTCAATTTATTTAGCACCAAAGACTCAAGCGCTGCGTGGGATGAGACCTTGTTGGATAAGTTTTACACCGAGTTATATCAACAACTCAACGATTTAGAGGCGTGTGTTATCCAAGGGGTCGGTGTGACGGAGACACCCTTGATGAAAGAGGACTCCATTCTTGCAGTGCGTAAGTACTTTCAACGCATTACGCTGTATCTTAAAGAGAAGAAGTACTCGCCGTGCGCATGGGAAGTCGTACGCGCAGAGATCATGCGCTCATTCTCGCTTTCAACGAACTTGCAAGAAAGCTTACGTAGTAAAGAG

1.12. **DARPin (528 bp): (7xHis-linker-cleavage site-DARPin)**

atgcatcatcatcatcatcatcacGGCGGTAGCGAAGCAGTGTATCATCAAGGTTCGGACTTAGGCCGCAAGTTACTGGAAGCTGCACGCGCGGGTCAAGACGACGAGGTACGCATCCTTATGGCCAATGGTGCGGACGTCAACGCCGCCGATAATACCGGAACAACTCCTTTGCACCTCGCAGCATATTCCGGCCATTTAGAGATTGTAGAGGTACTGCTTAAGCACGGTGCCGATGTAGATGCGTCGGACGTATTCGGGTACACGCCTCTCCACCTCGCCGCTTATTGGGGCCACTTAGAAATTGTAGAGGTGCTCTTGAAGAATGGTGCCGACGTGAATGCTATGGATTCAGACGGCATGACACCGCTCCATCTGGCCGCGAAATGGGGTTATCTGGAGATCGTCGAGGTTCTCTTGAAACATGGCGCCGACGTCAATGCACAAGATAAGTTCGGCAAGACTGCCTTCGACATCAGCATCGACAACGGTAATGAGGACCTGGCCGAGATCCTTCAAAAGTTGAAT

1.13. **GFP (1182 bp): (6xHis-linker-cleavage site-GFP-TRAP)**

atgcatcaccatcaccatcacGATTACGATATCCCAACGACCGAAGCAGTGTATCATCAAATGCGTAAAGGCGAAGAACTGTTCACGGGCGTAGTTCCGATTCTGGTCGAGCTGGACGGCGATGTGAACGGTCATAAGTTTAGCGTTCGCGGTGAAGGTGAGGGCGACGCGACCAACGGCAAACTGACCCTGAAGTTCATCTGCACCACCGGTAAACTGCCGGTGCCTTGGCCGACCTTGGTGACGACGTTGACGTATGGCGTGCAGTGTTTTGCGCGTTATCCGGACCACATGAAACAACACGATTTCTTCAAATCTGCGATGCCGGAGGGTTACGTCCAGGAGCGTACCATTTCCTTCAAGGATGATGGCTACTACAAAACTCGCGCAGAGGTTAAGTTTGAAGGTGACACGCTGGTCAATCGTATCGAATTGAAGGGTATCGACTTTAAAGAGGATGGTAACATTCTGGGCCATAAACTGGAGTATAACTTCAACAGCCATAATGTTTACATTACGGCAGACAAGCAAAAGAACGGCATCAAGGCCAATTTCAAGATTCGCCACAATGTTGAGGACGGTAGCGTCCAACTGGCCGACCATTACCAGCAGAACACCCCAATTGGTGACGGTCCGGTTTTGCTGCCGGATAATCACTATCTGAGCACCCAAAGCGTGCTGAGCAAAGATCCGAACGAAAAACGTGATCACATGGTCCTGCTGGAATTTGTGACCGCTGCGGGCATCACCCACGGTATGGACGAGCTGTATAAAGCCGGTGGTGGTTCTGGTGGTGGTTCGAAGCAGGCACTGAAAGAAAAAGAGCTGGGGAACGATGCCTACAAGAAGAAAGACTTTGACACAGCCTTGAAGCATTACGACAAAGCCAAGGAGCTGGACCCCACTAACATGACTTACATTATCAATCAAGCAGCGGTATACTTTGAAAAGGGCGACTACAATAAGTGCCGGGAGCTTTGTGAGAAGGCCATTGAAGTGGGGAGAGAAAACCGAGAGGACTATCGATGGATTGCCATTGCATATGCTCGAATTGGCAACTCCTACTTCAAAGAAGAAAAGTACAAGGATGCCATCCATTTCTATAACAAGTCTCTGGCAGAGCACCGAACCCCAAAGGTGCTAAAAAAGTGCCAACAGGCGGAGAAAATCCTGAAGGAGCAA

**2- Supplementary tables:**

| **2.1. Supplementary Table 1. Oligonucleotides (IDT, Belgium).** | |
| --- | --- |
| **2.1.1. Primers for Con1 fragmentation:** | |
| FOR: | AACAAGGACCATAGATTATGCATCATCATCATCATCATCAC |
| REV: | ACTCTAGAGGATCCTTATTAGTCCGTTACTAATTTTACGA |
| **2.1.2. Primers for pMAL linearization for Con1:** | |
| FOR: | TCGTAAAATTAGTAACGGACTAATAAGGATCCTCTAGAGTC |
| REV: | TGATGATGATGATGATGATGCATAATCTATGGTCCTTGTT |
| **2.1.3. Primers for TuMV_J fragmentation:** | |
| FOR: | AACAAGGACCATAGATTATGCATCATCATCATCATCATCAC |
| REV: | ACTCTAGAGGATCCTTATTAATCTAAGTCGGAAATAAGTTT |
| **2.1.4. Primers for pMAL linearization for TuMV_J:** | |
| FOR: | AACTTATTTCCGACTTAGATTAATAAGGATCCTCTAGAGTC |
| REV: | TGATGATGATGATGATGATGCATAATCTATGGTCCTTGTT |
| **2.1.5. Primers for Stefin A fragmentation:** | |
| FOR: | CATCAAATGATTCCCGGCGGTTTATCAGAGGC |
| REV: | AGGTCGACTCTAGAGGATCCTTAAAATCCAGTGAGTTCGTCG |
| **2.1.6. Primers for pMAL linearization for Stefin A:** | |
| FOR: | ACGAACTCACTGGATTTTAAGGATCCTCTAGAGTCGACC |
| REV: | GGGAATCATTTGATGATACACTGCTTCGCTACCG |
| **2.1.7. Primers for Interferon α2 fragmentation:** | |
| FOR: | AGCGAAGCAGTGTATCATCAAATGTGCGACTTGCCCCAGAC |
| REV: | GACTCTAGAGGATCCTTACTCTTTACTACGTAAGCTTTCTTGCAAG |
| **2.1.8. Primers for pMAL linearization for Interferon α2:** | |
| FOR: | GAAAGCTTACGTAGTAAAGAGTAAGGATCCTCTAGAGTCGACC |
| REV: | CTGGGGCAAGTCGCACATTTGATGATACACTGCTTCGCTAC |
| **2.1.9. Primers for DARPin fragmentation:** | |
| FOR: | GCGAAGCAGTGTATCATCAAGGTTCGGACTTAGGCCG |
| REV: | ACTCTAGAGGATCCTTAATTCAACTTTTGAAGGATCTCGG |
| **2.1.10. Primers for pMAL linearization for DARPin:** | |
| FOR: | TCCTTCAAAAGTTGAATTAAGGATCCTCTAGAGTCGACC |
| REV: | TTGCGGCCTAAGTCCGAACCTTGATGATACACTGCTTCGC |
| **2.1.11. Primers for mutating TEV site to Con1 site in His-TEV-GFP:** | |
| FOR: | GCAGTGTATCATCAAATGCGTAAAGGCGAAGAACTG |
| REV: | TTGATGATACACTGCTTCGGTCGTTGGGATATCGTAAT |
| **2.1.12. Primers for mutating the first residue of DARPins (the mutant residue is underlined)** | |
| **A**-Dar.FOR: | TATCATCAAGCTTCGGACTTAGGCCGC |
| **A**-Dar.REV: | CGAAGCTTGATGATACACTGCTTCGCTAC |
| **E**-Dar.FOR: | TATCATCAAGAATCGGACTTAGGCCGC |
| **E**-Dar.REV: | CGATTCTTGATGATACACTGCTTCGCTAC |
| **F**-Dar.FOR: | GTATCATCAATTTTCGGACTTAGGCCGC |
| **F**-Dar.REV: | CGAAAATTGATGATACACTGCTTCGCTACC |
| **R**-Dar.FOR: | TATCATCAACGTTCGGACTTAGGCCGC |
| **R**-Dar.REV: | CGAACGTTGATGATACACTGCTTCGCTAC |
| **S**-Dar.FOR: | TATCATCAATCCTCGGACTTAGGCCGC |
| **S**-Dar.REV: | CGAGGATTGATGATACACTGCTTCGCTAC |
| **T**-Dar.FOR: | TATCATCAAACCTCGGACTTAGGCCGC |
| **T**-Dar.REV: | CGAGGTTTGATGATACACTGCTTCGCTAC |
| **Q**-Dar.FOR: | TATCATCAACAGTCGGACTTAGGCCGC |
| **Q**-Dar.REV: | CGACTGTTGATGATACACTGCTTCGCTAC |
| **W**-Dar.FOR: | TATCATCAATGGTCGGACTTAGGCCGC |
| **W**-Dar.REV: | CGACCATTGATGATACACTGCTTCGCTAC |
| **M-**Dar.FOR: | TATCATCAAATGTCGGACTTAGGCCGC |
| **M-**Dar.REV: | CGACATTTGATGATACACTGCTTCGCTAC |
| **D**-Dar.FOR: | TATCATCAAGATTCGGACTTAGGCCGC |
| **D**-Dar.REV: | CGAATCTTGATGATACACTGCTTCGCTAC |
| **I**-Dar.FOR: | TATCATCAAATCTCGGACTTAGGCCGC |
| **I**-Dar.REV: | CGAGATTTGATGATACACTGCTTCGCTAC |
| **2.1.13. Primers for deleting 228-234 residues of Con1** | |
| FOR: | GAGCCTTTCAAATAATAAGGATCCTCTAGAGTC |
| REV: | TTATTTGAAAGGCTCTTCTGGTTGGCTTTC |
| **2.1.14. Primers for deleting 222-234 residues of Con1** | |
| FOR: | AAGAAAGCCAATAATAAGGATCCTCTAGAGTC |
| REV: | TTATTGGCTTTCTTGAAGGTTAAGGCTCC |

| **2.2. Supplementary Table 2. Predicted solubility profiles of the selected proteases.** | |
| --- | --- |
| **Protease** | **Intrinsic solubility score** |
| TEV | 0.47 |
| PVA | -0.03 |
| TuMV_Q | 0.64 |
| TuMV_J | 0.71 |
| PPV | 0.57 |
| OMV | 0.65 |
| TVMV | 0.45 |
| LMVE | 0.31 |
| LMV0 | 0.30 |
| Sequence-based solubility calculation by CamSol. | |

| **2.3. Supplementary Table 3. Designed consensus amino acid sequences.** | |
| --- | --- |
| **Design** | **Sequence** |
| Con1 | SKSLFRGLRDYNPIASNICHLTNESDGHSNSLYGIGFGPLIITNQHLFRRNNGELTIQSRHGEFVVKNTTQLKLLPIDGRDILIIRLPKDFPPFPQKLKFRQPEKGERICLVGSNFQTKSITSTVSETSTTMPVENSQFWKHWISTKDGHCGSPLVSTKDGKILGILSLSNFTNTNNYFAAFPEDFEETYLHTQEAQEWVKHWKYNPDAISWGSLNLQESLPEEPFKIVKLVTDLFSDAVYAQ |
| Con2 | SKSLFRGLRDYNPISNAICHLTNESDGHSESLYGIGFGPLIITNQHLFRRNNGELRVQSRHGEFVVKNTTQLKLLPCEGRDIIVIRLPKDFPPFPQKLKFRQPIKGERICLVGSNFQTKSISSTVSETSVTTPVDNSFFWKHWISTKDGQCGLPLVSTKDGFIVGIHSLTNSTNTQNYFAAFPDDFHETYLADIEAHSWVKHWKYNPDEVSWGGLNLQASTPREPFKIIKLVTDLDGDAVYFQ |
| Con3 | SKSLFRGLRDYNPISSAICHLTNESDGHSESLYGIGFGPLIITNQHLFRRNNGELTVQSRHGEFVVKNTTQLKLLPIEGRDILVIRLPKDFPPFPQKLKFRQPEKGERICLVGSNFQTKSITSTVSETSVTMPVENSQFWKHWISTKDGHCGSPLVSTKDGKILGIHSLANFTNTQNYFAAFPEDFEETYLHTQEAQEWVKHWKYNPDAVSWGSLNLQESQPEEPFKIVKLVTDLFSDAVYAQ |
| Con4 | SKSLFRGLRDYNPISNNICHLTNESDGHSESLYGIGFGPLIITNQHLFRRNNGELRIQSRHGEFVVKNTTQLKLLPCEGRDIIIIRLPKDFPPFPQKLKFRQPIKGERICLVGSNFQTKSISSTVSETSTTTPVDNSFFWKHWISTKDGQCGLPLVSTKDGFIVGIHSLTNSTNTQNYFAAFPDDFHETYLADIEAHSWVKHWKYNPDEISWGGLNLQASTPREPFKIIKLVTDLDGDAVYFQ |
| Con5 | SKSLFRGLRDYNPISSAICHLTNESDGHSESLYGIGFGPLIITNQHLFRRNNGELTVQSRHGEFVVKNTTQLKLLPIEGRDILVIRLPKDFPPFPQKLKFRQPEKGERICLVGSNFQTKSITSTVSETSVTMPVENSQFWKHWISTKDGHCGSPLVSTKDGKILGIHSLANFTNTQNYFAAFPEDFEETYLADQEAHEWVKHWKYNPDAVSWGSLNLQESQPEEPFKIVKLVTDLFSDAVYAQ |
| Con6 | SKSLFRGLRDYNPISNNICHLTNESDGHSESLYGIGFGPLIITNQHLFRRNNGELRIQSRHGEFVVKNTTQLKLLPCEGRDIIIIRLPKDFPPFPQKLKFRQPIKGERICLVGSNFQTKSISSTVSETSTTTPVDNSFFWKHWISTKDGQCGLPLVSTKDGFIVGIHSLTNSTNTQNYFAAFPDDFHETYLHTIEAQSWVKHWKYNPDEISWGGLNLQASTPREPFKIIKLVTDLDGDAVYFQ |
| Con7 | SKSLFRGLRDYNPISNAICHLTNESDGHSESLYGIGFGPLIITNQHLFRRNNGELTVQSRHGEFVVKNTTQLKLLPIEGRDIILIRLPKDFPPFPQKLKFRQPEKGERICLVGSNFQTKSISSTVSETSTTMPVENSQFWKHWISTKDGQCGSPLVSTKDGFIVGIHSLSNFTNTQNYFAAFPDDFAETYLHTIEAQSWVKHWKYNPDAVSWGGLNLQASKPEEPFKISKLVTDLDGDAVYAQ |
| Con8 | SKSLFRGLRDYNPISSNICHLTNVSDGHSNSLYGIGFGPLIITNQHLFRRNNGELTIKSRHGEFVVKNTTQLHLLPIPGRDIIIIRLPKDFPPFPQKLKFRQPEKGERICMVGSNFQTKSISSMVSETSTIMPVENSQFWKHWISTKDGQCGSPLVSTKDGKILGLHSLANFTNTINYFAAFPDDFAETYLHTQEADEWVKHWKYNTDAISWGSLNLQASQPESPFKVSKLITDLDSTAVYAQ |
| Con9 | SNSLFRGLRDYNPISNNICHLTNVSDGASNSLYGVGFGPLILTNRHLFERNNGELTIKSRHGEFVVKNTTQLQLLPIPGRDLILIRLPKDFPPFPQKLGFRQPTKEERICMVGSNFQTKSVSSVVSETSTIMPVENSQFWKHWISTKDGQCGSPMVSTRDGKILGLHSLANFTNSINYFAAFPDNFMEKYLHNIEAHEWVKHWKYNADAISWGSLNIQASQPESLFKVSKLISDLDSELVYAQ |
| Con10 | SKSLFRGLRDYNPISNNICHLTNVSDGHSNSLYGIGFGPLIITNRHLFRRNNGELTIKSRHGEFVVKNTTQLHLLPIPGRDIIIIRLPKDFPPFPQKLKFRQPEKGERICMVGSNFQTKSISSVVSETSTIMPVENSQFWKHWISTKDGQCGSPLVSTKDGKILGLHSLANFTNTINYFAAFPDDFMETYLHTIEADEWVKHWKYNTDAISWGSLNLQASQPESPFKVSKLITDLDSTAVYAQ |
| C-terminal (235-243) shown in blue. | |

**3- Supplementary figures:**

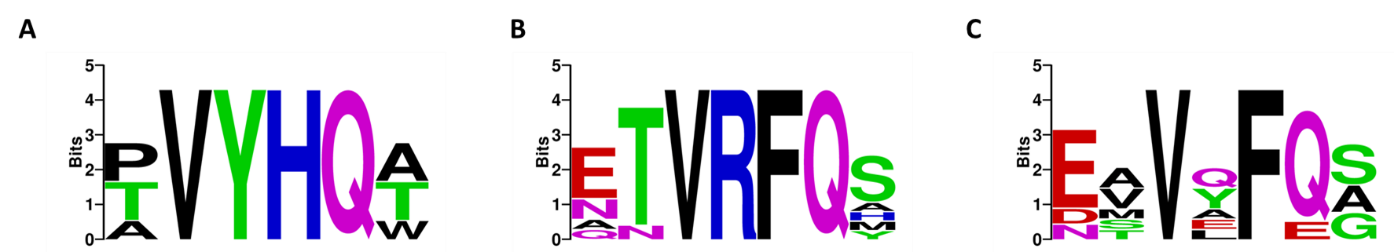

**Supplementary Figure 1.** The conserved sequence of the cleavage site of TuMV_Q or PPV proteases; n=5 (**A**), TVMV protease; n=9 (**B**) or PVA protease; n=5 (**C**). The cleavage sites were identified from the MEROPS database, and the logo plots were generated by the [WebLogo](https://weblogo.berkeley.edu/logo.cgi) online application.

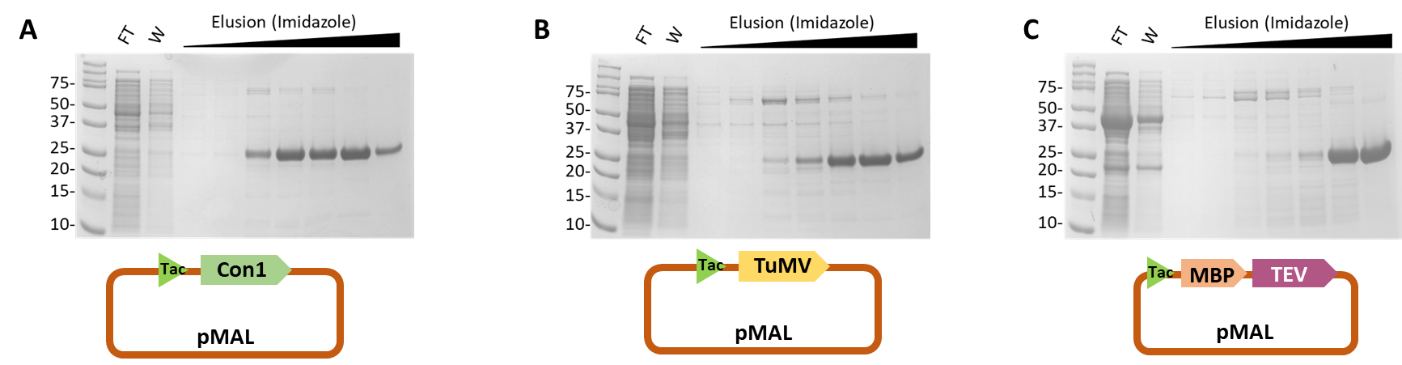

**Supplementary Figure 2**. SDS-PAGEs showing the purification steps of N-terminal His tagged proteases. (**A** - **C**) Gel photos showing the purification of Con1 (**A**), TuMV_J (**B**) or TEV (**C**) that have been expressed by pMAL vector and Tac promoter. The TEV plasmid (pRK793; Addgene #8827) contains maltose binding protein (MBP) upstream to TEV gene. The purification was done with 5-ml Ni-Sepharose resin in home-packed columns. The molecular weights are around 27 kDa for Con1 and TuMV_J, and 28.6 for TEV. FT: flow through; W: wash.

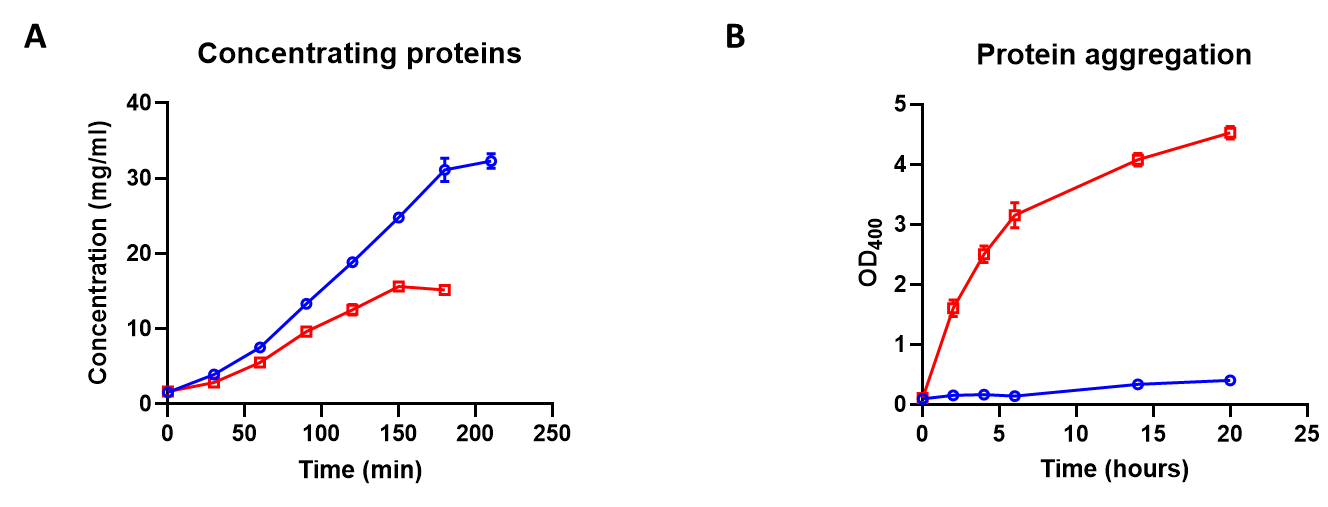

**Supplementary Figure 3.** (**A**) TEV (red) or Con1 (blue) proteases concentration over a 3K MWCO Amicon concentrator. Aliquots were collected every 30 min and the concentrations were measured by Nano-drop spectrophotometer in duplicate readings using the extension coefficient of 31970 and 34950 M^-1^cm^-1^ for TEV (28560.45 Da) and Con1 (27660.22 Da), respectively. (**B**) The optical density tracking of 15 mg/ml TEV and 30 mg/ml Con1 at 400 nm, measured by Nano-drop in duplicate readings.

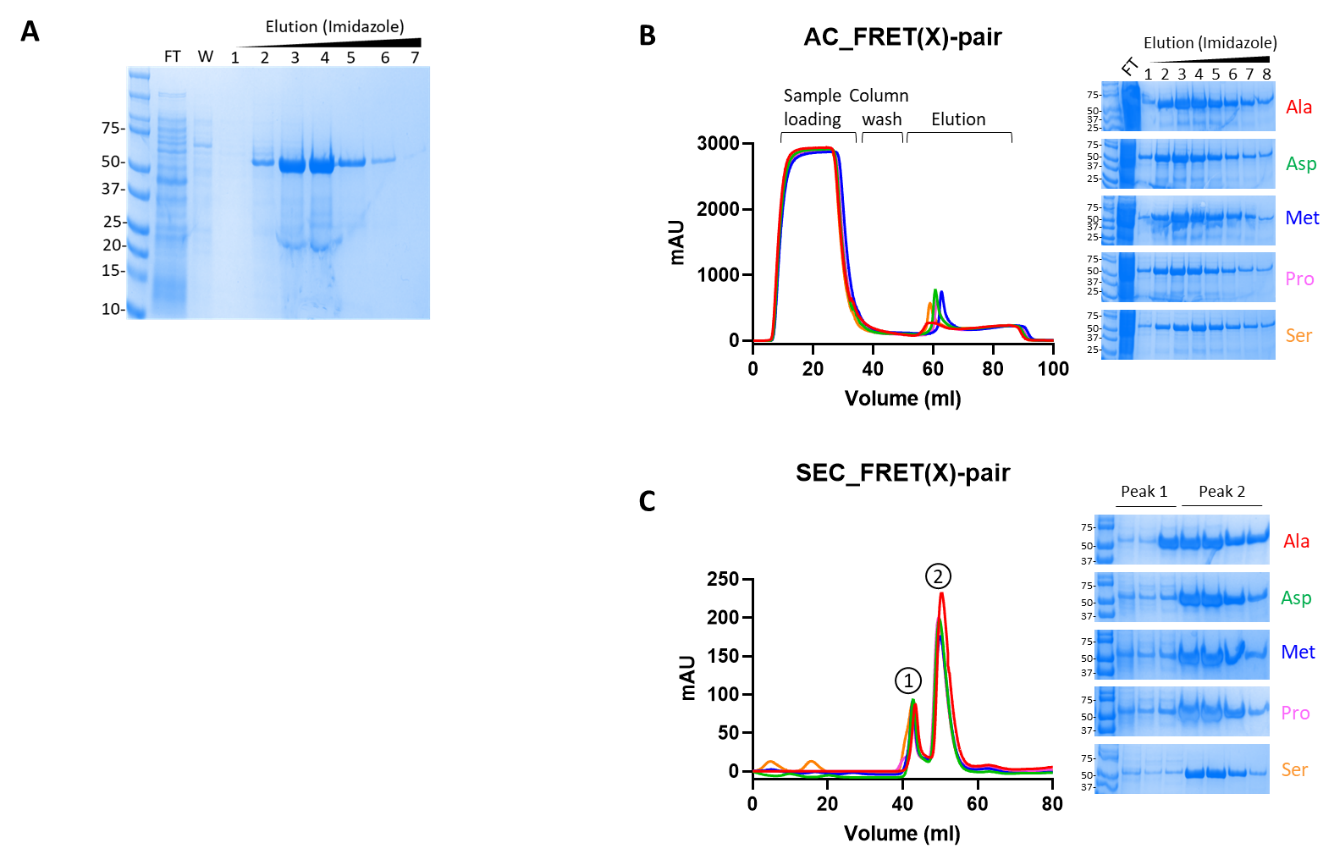

**Supplementary Figure 4**. Purification of FRET(X) substrates (CFP-cleavage site-YFP) with different P1’ (X). (**A**) SDS-PAGE photo showing the purification of FRET-ENLYFQG substrate using 5-ml Ni-Sepharose resin in home-packed column. (**B**) Affinity chromatography (AC) using HisTrap excel Ni column showing the eluted fractions on the SDS-PAGE. (**C**) Size exclusion chromatography (SEC) over Superdex 75 column showing the excluded peaks on SDS-PAGE. The purification steps were done by AKTA Go instrument. X = alanine, serine, aspartic, methionine or proline. The molecular weights are around 60 kDa as theoretically calculated. FT: flow through; W: wash.

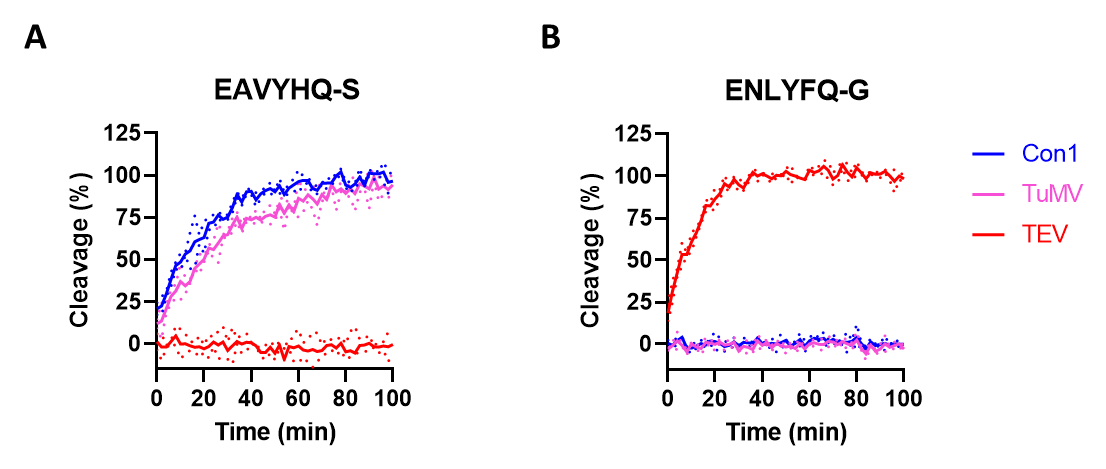

**Supplementary Figure 5.** Time-course screening of Con1 protease specificity against different FRET substrates. (**A** and **B**) Screening Con1, TuMV_J or TEV specificity against the different cleavage sites, EAVYHQS (**A**) or ENLYFQG (**B**), introduced to FRET substrates. The reaction composed of 1 µM enzyme and 2 µM substrate final concentrations and the fluorescence was measured over 2 hours period at room temperature by the plate reader. Cleavage is a percent of maximum. FRET ratio was calculated by dividing the fluorescence intensity at 480 nm (donor) by that at 520 nm (acceptor), blank subtraction and percentage calculation. All the results are the mean of 2 technical repeats ± standard deviation.

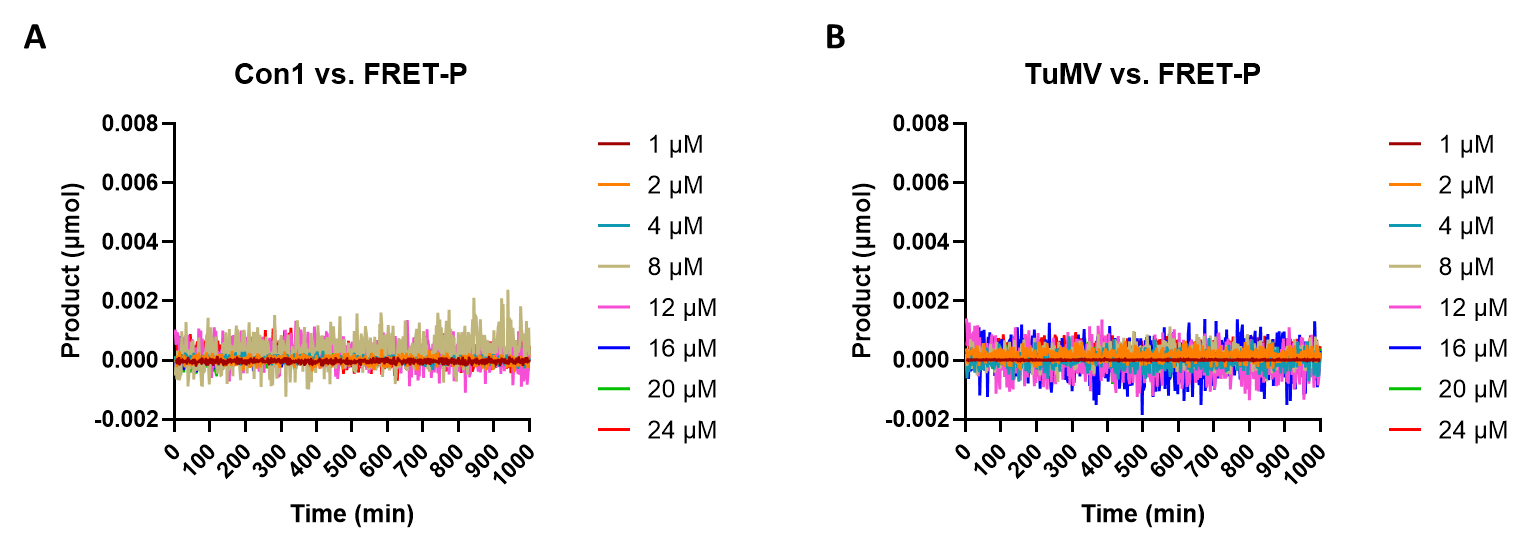

**Supplementary Figure 6**. Time-course screening specificity of Con1 (**A**) and TuMV_J (**B**) against EAVYHQ-P. The reaction mixture contains 0.4 µM protease with the indicated substrate concentration and the fluorescence was gained over 17 hours period at room temperature by the plate reader. The results are the mean of 2 technical repeats (solid lines) ± standard deviation (dotted lines). The curves were fitted by the exponential plateau equation on GraphPad Prism (black lines) and the reaction rates were calculated.

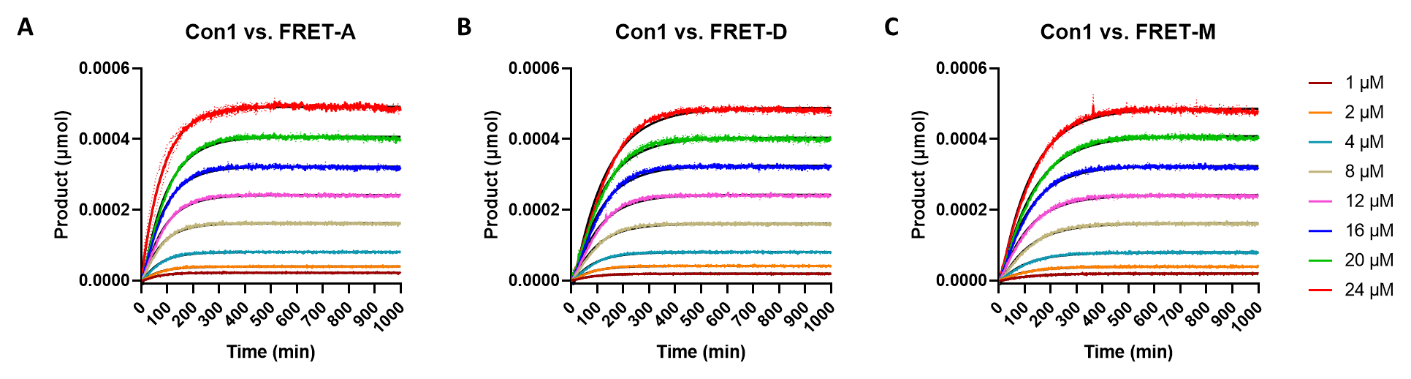

**Supplementary Figure 7**. Kinetic studies of Con1 protease activity against different P_1_’ FRET substrates where P_1_’ = alanine (**A**), aspartic (**B**), or methionine (**C**). The reaction mixture contains 0.4 µM Con1 with the indicated substrate concentration and the fluorescence was gained over 17 hours period at room temperature by the plate reader. The results are the mean of 2 technical repeats (solid lines) ± standard deviation (dotted lines). The curves were fitted by the exponential plateau equation on GraphPad Prism (black lines) and the reaction rates were calculated.

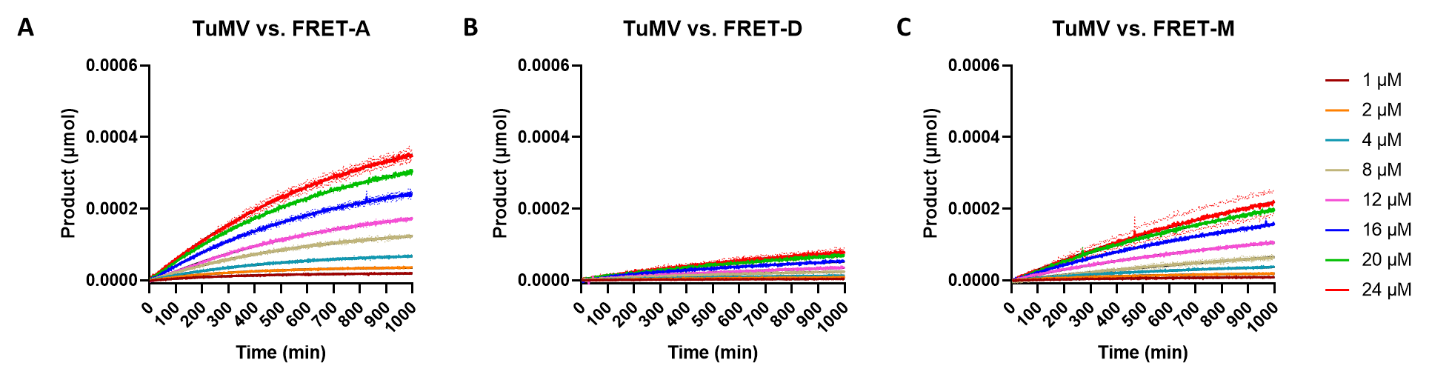

**Supplementary Figure 8**. Kinetic studies of TuMV_J protease activity against different P_1_’ FRET substrates where P_1_’ = alanine (**A**), aspartic (**B**), or methionine (**C**). The reaction mixture contains 0.4 µM TuMV_J with the indicated substrate concentration and the fluorescence was gained over 17 hours period at room temperature by the plate reader. The results are the mean of 2 technical repeats (solid lines) ± standard deviation (dotted lines). The curves were fitted by the exponential plateau equation on GraphPad Prism (black lines) and the reaction rates were calculated.

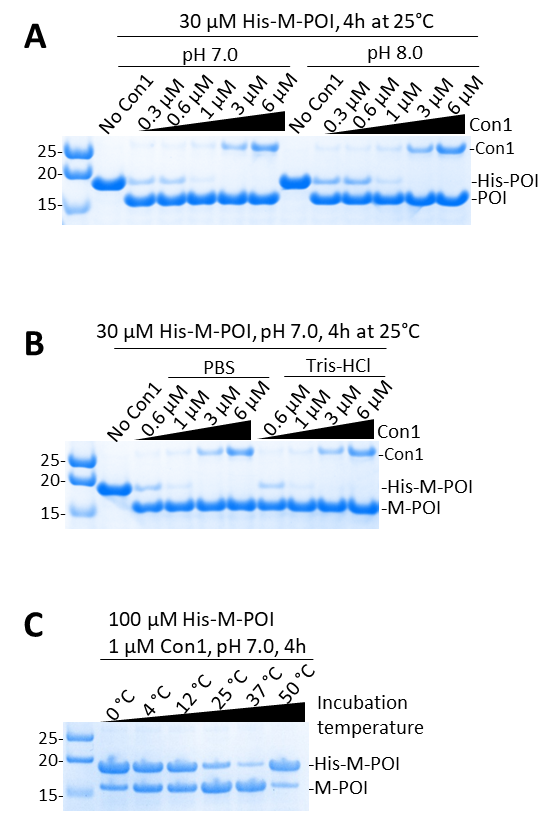

**Supplementary Figure 9.** Characterization of the pH and temperature dependency of Con1 activity. (**A** and **B**) Study the effect of different pH (**A**; 7.0 or 8.0 of Tris-HCl buffer) or different buffers (**B**) on the Con1 activity to cleave the His tag at different ratios. (**C**) Study the effect of increasing the reaction temperature on the Con1 activity. Con1 and substrate were pre-incubated at the indicated temperatures.

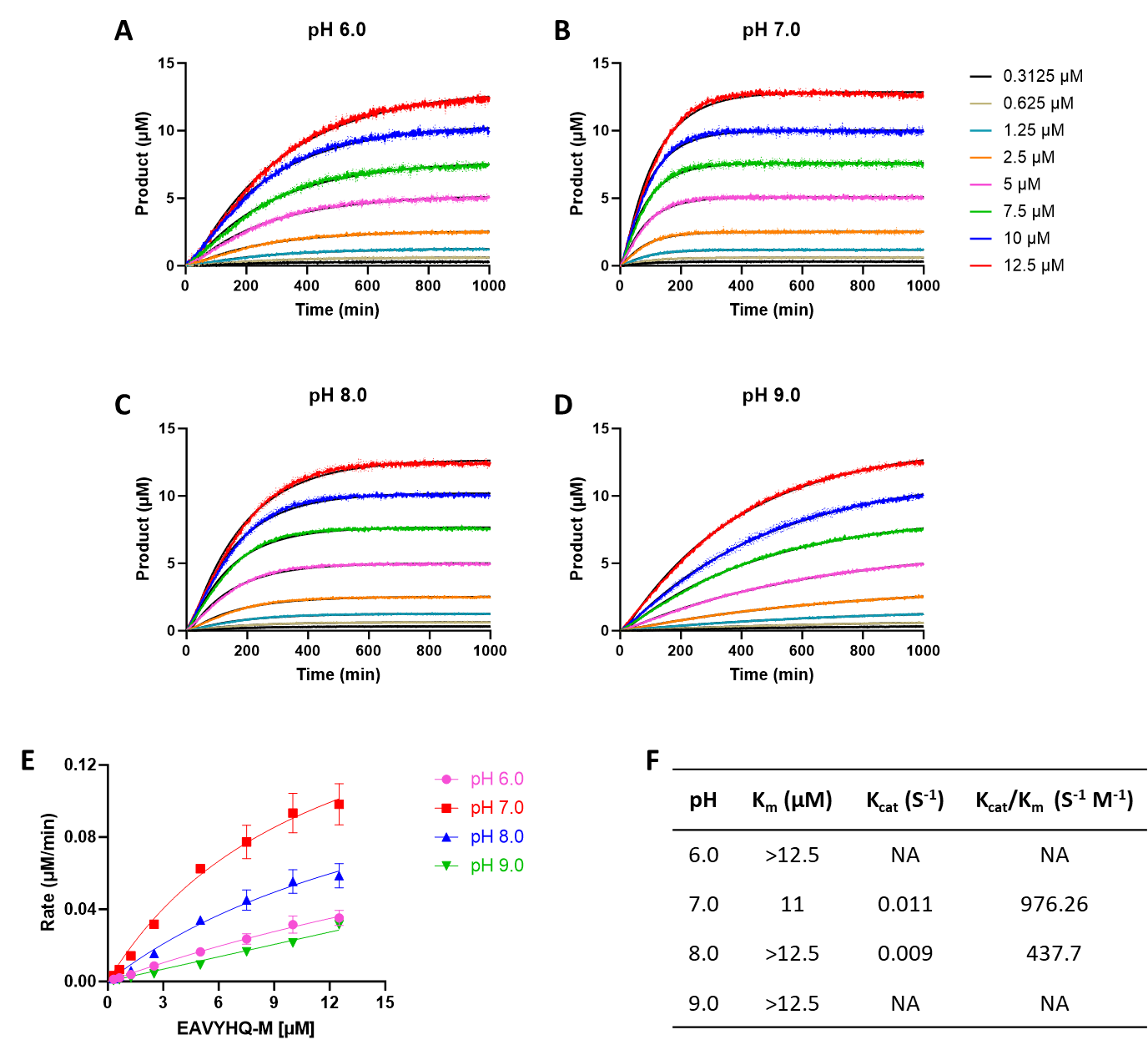

**Supplementary Figure 10.** Kinetic studies of Con1 protease activity against FRET-M substrate at different pH as indicated. Con1 and substrate fractions were pre-incubated at different pH and the reaction mixture contained 0.4 µM Con1 with the indicated substrate concentrations at (A) 0.1M citrate, pH 6.0; (B) 0.1M Tris-HCl, pH 7.0; (C) 0.1M Tris-HCl, pH 8.0 or (D) 0.1M Tris-HCl, pH 9.0). The fluorescence was gained over 17-hour period at room temperature by the plate reader. The results are the mean of 2 technical repeats (solid lines) ± standard deviation (dotted lines). The curves were fitted by the exponential plateau equation on GraphPad Prism (black lines) and the initial rates were calculated. (E) Kinetic summary of Con1 at the different pH. The results are the mean of 2 technical repeats ± standard deviation. The curves are fitted by the Michaelis-Menten non-linear regression using GraphPad Prism and the kinetic parameters are shown in (F).

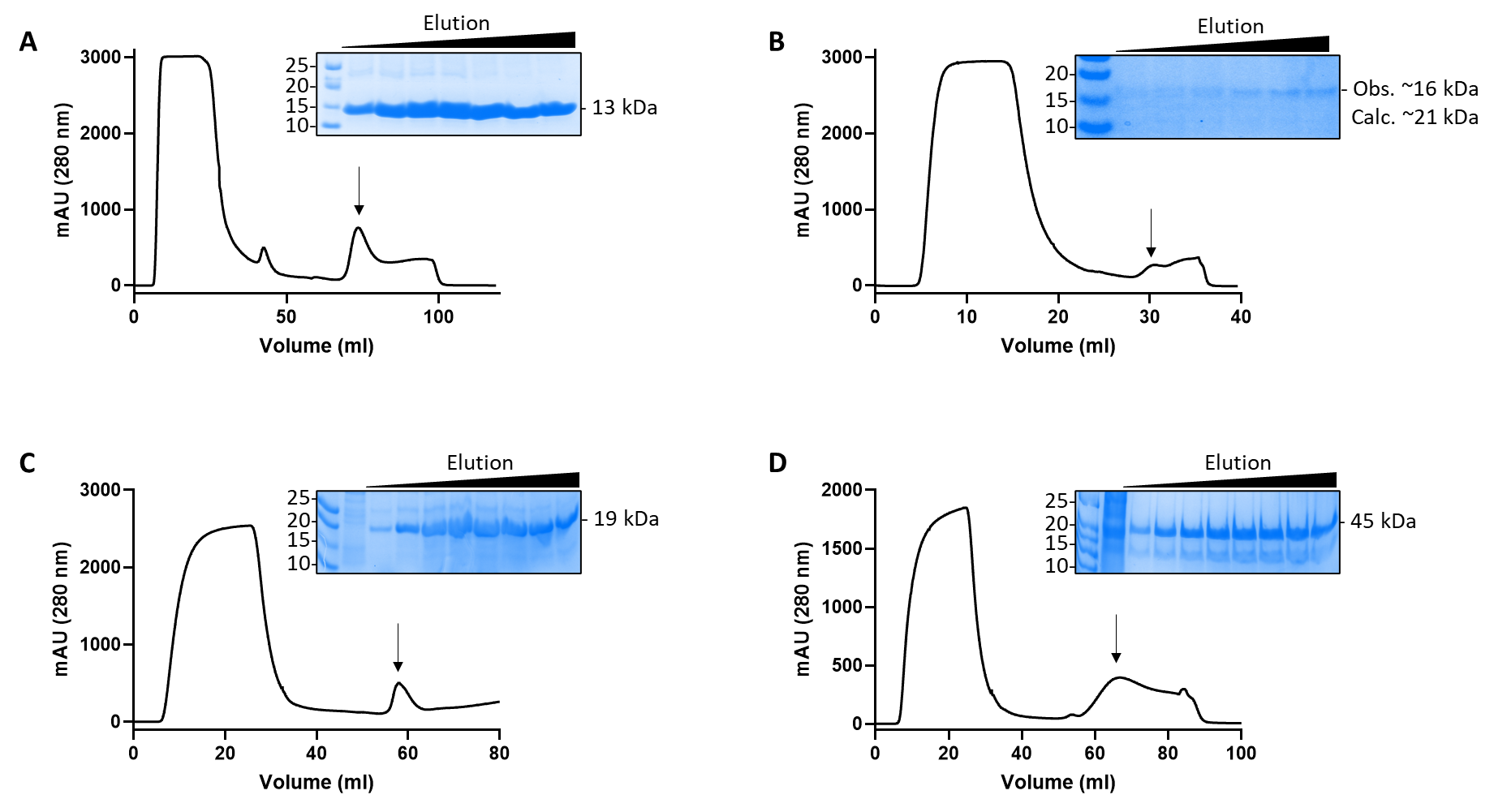

**Supplementary Figure 11.** (**A**-**C**) Purification of His-tagged proteins of interest (POI) of StefinA (**A**), Interferon α2 (**B**), DARPins (**C**) and GFP (**D**) with Con1 cleavage site between the His tag and the POI. The purification was done by one-step affinity chromatography (AC) using HisTrap excel Ni column on an AKTA Go instrument and the eluted fractions are shown on the SDS-PAGEs.

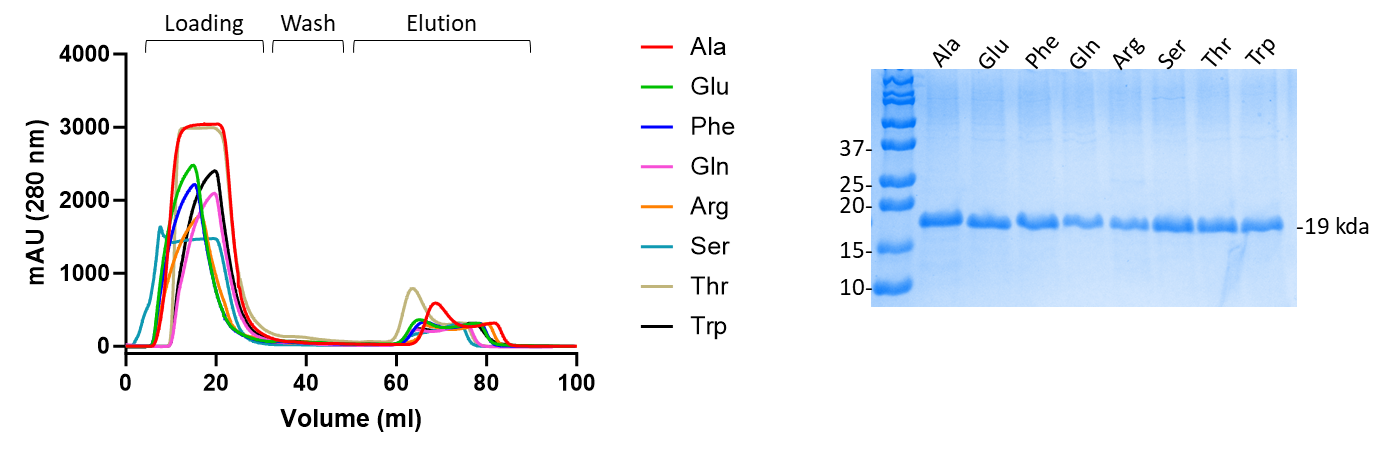

**Supplementary Figure 12.** Purification of His-tagged DARPins with different N-terminal as indicated with Con1 cleavage site between the His tag and the POI. The purification was done by one step affinity chromatography (AC) using HisTrap excel Ni column on an AKTA Go instrument and the eluted fractions are shown on the SDS-PAGE.

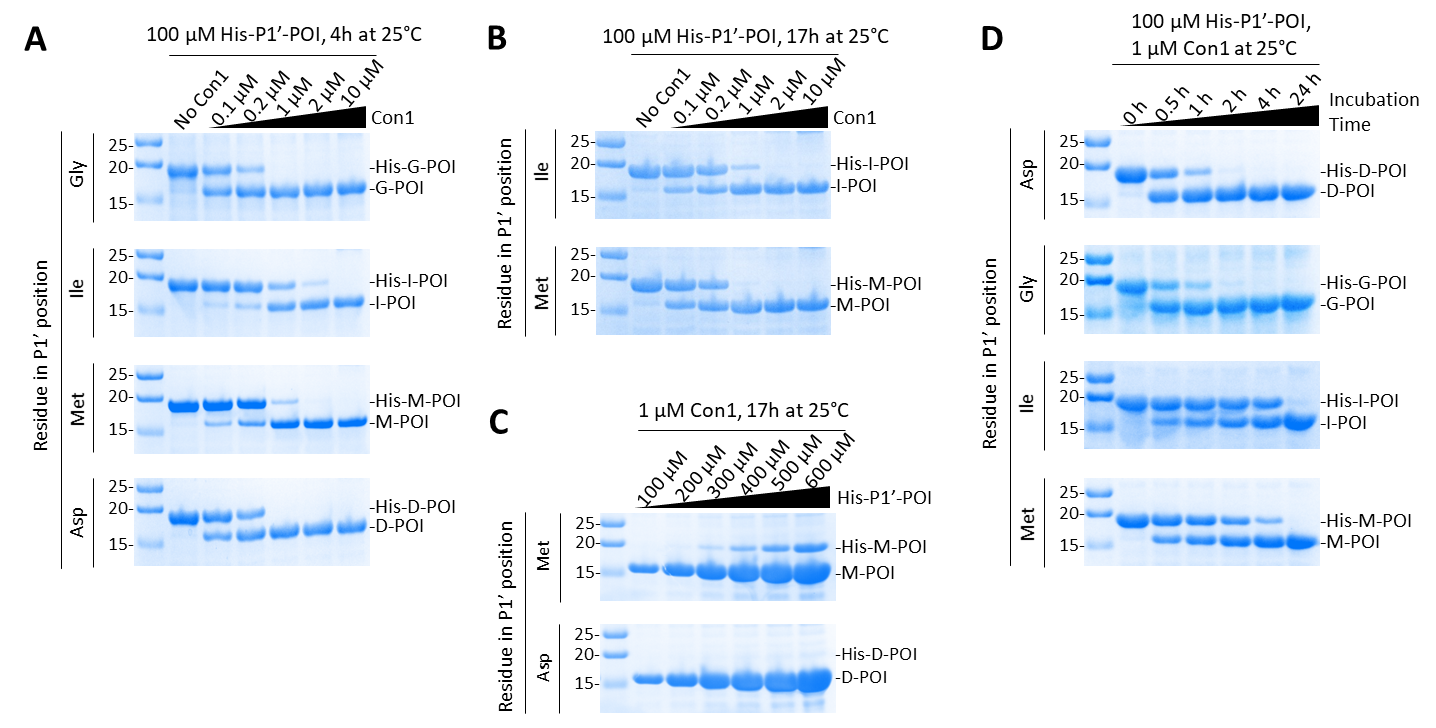

**Supplementary Figure 13.** SDS-PAGE experiments showing the His tag removal by Con1. (A-D) Study the enzyme: substrate ratio variation against the incubation time required to fully cleave the His tag from different P1’ positions representing the different N-terminal of POI. (A and B) Increasing Con1 concentration against 100 µM of different substrates as indicated that was incubated for 4h (A) or 17h (B) at room temperature. (C) Increasing substrate concentration against 1 µM Con1 that was incubated for 17h at room temperature. (D) Study the incubation time required to fully remove the His tag from different substrates as indicated at an enzyme: substrate molar ratio of 1: 100 at room temperature.

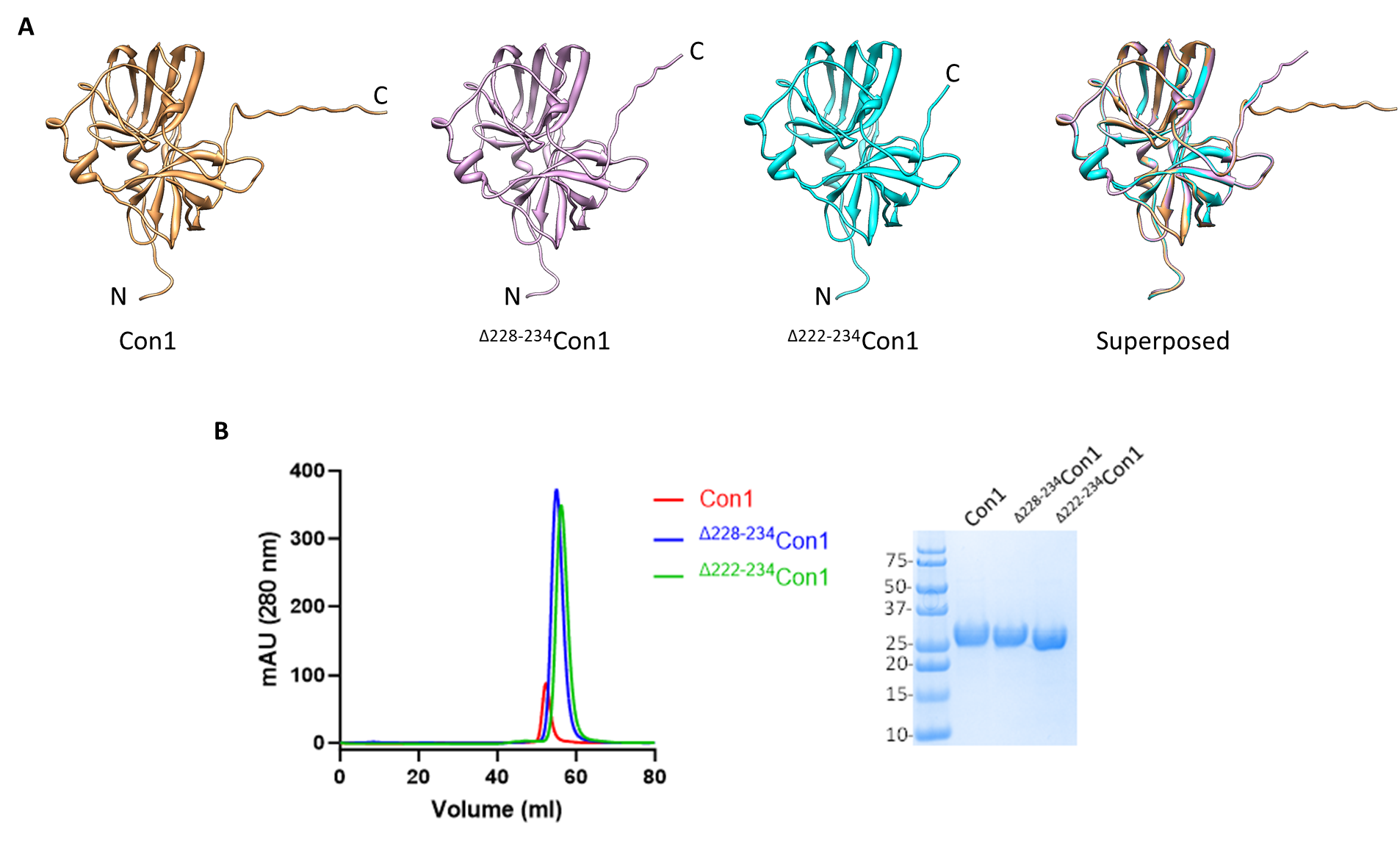

**Supplementary Figure 14.** Preparation of C-terminal truncated versions of Con1. (**A**) Ribbon representations of the AlphaFold2 predicted models showing the unstructured C-terminal of Con1 (221-234) and the shorter versions by deleting 228-234 (^Δ228-234^Con1) or 222-234 (^Δ222-234^Con1) residues. (**B**) Purification of His-tagged Con1 versions as indicated. The purification was done by one step affinity chromatography (AC) using HisTrap excel Ni column on an AKTA Go instrument and the eluted fractions are shown on the SDS-PAGE.

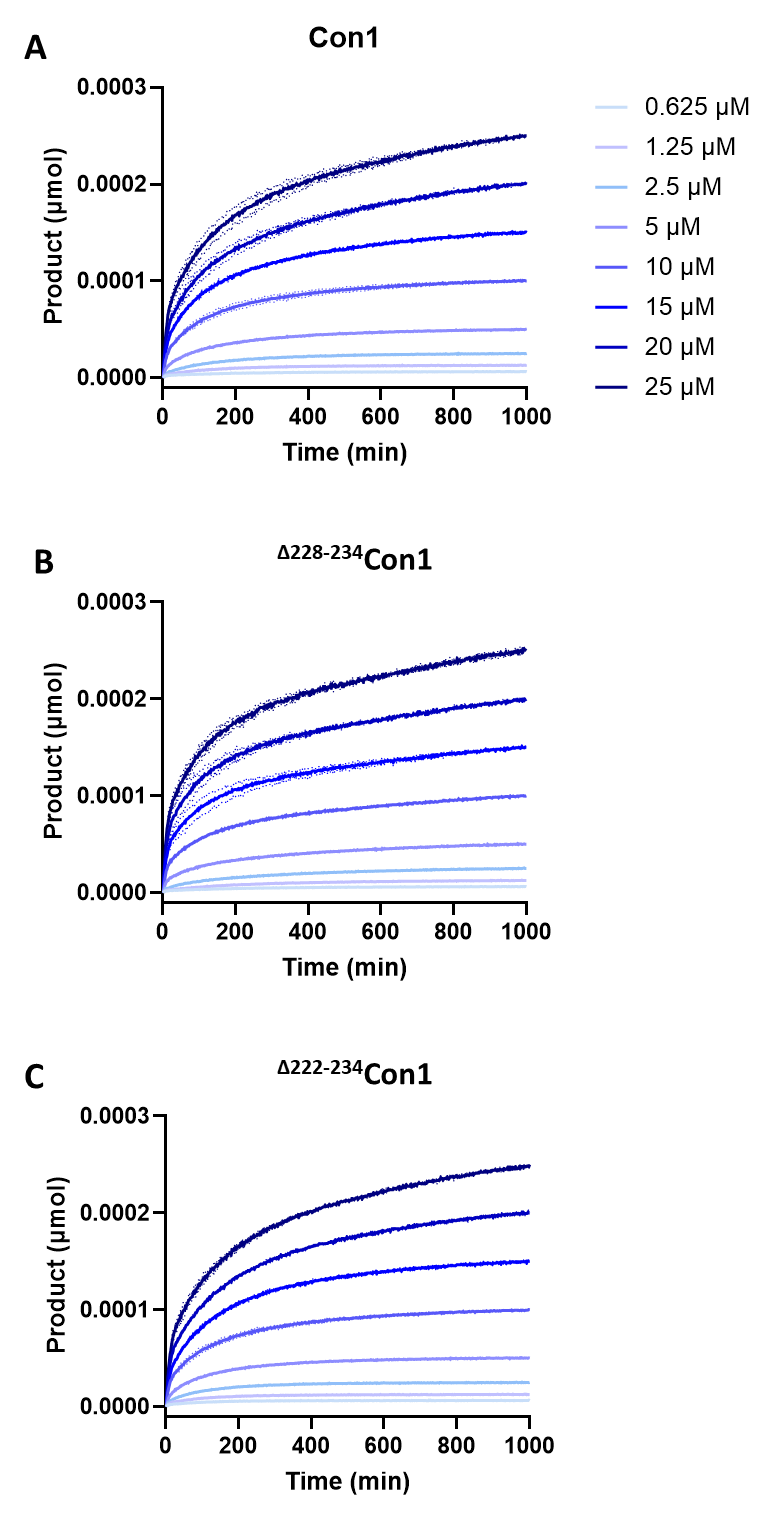

**Supplementary Figure 15.** Kinetic studies of the different Con1 versions as indicated against FRET-M substrate. The reaction mixture contains 0.4 µM protease with the indicated substrate concentration and the fluorescence was gained over 17 hours period at room temperature by the plate reader. The results are the mean of 2 technical repeats (solid lines) ± standard deviation (dotted lines). The curves were fitted by the exponential plateau equation on GraphPad Prism (black lines) and the initial rates were calculated.
